# Functional imaging and connectome analyses reveal organizing principles of taste circuits in *Drosophila*

**DOI:** 10.1101/2024.08.23.609242

**Authors:** Jinfang Li, Rabiah Dhaliwal, Molly Stanley, Pierre Junca, Michael D. Gordon

## Abstract

Taste is crucial for many innate and learned behaviors. In the fly, taste impacts feeding, oviposition, locomotion, mating, and memory formation, to name a few. These diverse roles may necessitate the apparent distributed nature of taste responses across different circuits in the fly brain, leading to complexity that has hindered attempts to deduce unifying principles of taste processing and coding. Here, we combine information from the whole brain connectome with functional calcium imaging to examine the neural representation of taste at early steps of processing. We find that the representation of taste quality remains largely segregated in cholinergic and GABAergic local interneurons (LNs) that are directly postsynaptic to taste sensory neurons of the labellum. Although some taste projection neurons (TPNs) projecting to superior protocerebrum receive direct inputs from sensory neurons, many receive primarily indirect taste inputs via cholinergic LNs. Moreover, we found that cholinergic interneurons appear to function as nodes to convey feedforward information to dedicated sets of morphologically similar TPNs. Examining a small number of representative TPNs suggests that taste information remains mostly segregated at this level as well. Together, these studies suggest a previously unappreciated logic in the organization of fly taste circuits.

## Introduction

Chemosensation plays an important role in guiding animal behaviors such as foraging, feeding, homeostasis, reproduction, and social activities (1–8). Olfaction and taste are the main chemical senses, and each involves translating environmental chemical stimuli into electrical activity that can be decoded by downstream neural circuits. However, the logic of taste coding remains controversial across multiple systems, from mammals to insects. Moreover, in contrast to olfaction, little is known about the global organization of taste circuits in the important model system *Drosophila melanogaster*.

In mammals, taste information is detected by sweet, bitter, appetitive salt, umami, and sour taste receptor cells (TRCs; 9–11). The detected taste information is sequentially relayed from cranial nerves to the nucleus of solitary tract (NST), parabronchial nucleus (PBN), parvicellular portion of the VPMpc, and insular cortex (10, 12). The relay stations at several points in the ascending pathway also pass on taste information to limbic systems for functions like multisensory integration and associative learning (12). Within the rostral NST, inhibitory local neurons modulate tuning width and enhance contrast between pleasant vs unpleasant tastes (13–17). However, the overall coding model for early processing of mammalian taste is subject to debate. For example, imaging studies on geniculate ganglion neurons show evidence of labelled line and combinatorial coding models, but the extent to which each model is applied varies (18, 19).

The *Drosophila* olfactory system has been a powerful model for understanding the functional organization and early processing of chemosensory information. Many odorant receptors are broadly tuned to various ligands, and olfactory receptor neurons (ORNs) expressing the same receptor converge on the same glomerulus of the antennal lobe, where they form synaptic connections to several olfactory projection neurons (PNs; 8, 20, 21). Within the antennal lobe, there are broad GABAergic local neurons (LNs) that receive signals from ORNs and mediate presynaptic inhibition of ORNs (8, 20–24). The activity of the inhibitory LNs is scaled based on the presynaptic input strength, thereby mediating input gain control (8, 20, 21, 23, 25–28). This presynaptic gain control has an important functional role in preventing the saturation of glomeruli and increasing decoding efficiency (25–28). Recently, a class of distinct LNs called ‘patchy’ cells was shown to be morphologically and functionally distinct from broad LNs and provide intra-glomerular output gain control of PN responses (29, 30). In addition to inhibitory LNs, cholinergic LNs in the antennal lobe form gap junctions with PNs, causing correlated activity in other glomeruli and expanding PN tuning (20, 31–33). Ultimately, olfactory information carried by PNs is sent to the mushroom body (MB) and lateral horn (LH) to mediate learned and innate olfactory behaviors, respectively (8, 21).

In contrast to the delineated gustatory pathways in mammals and olfactory circuits in flies, *Drosophila* does not have an accepted primary circuit for taste information (3, 10, 12). Rather, taste pathways appear widely distributed, making it challenging to understand the logic of taste information flow in the fly brain. Tastes are detected by gustatory receptor neurons (GRNs) distributed throughout the body including the labella, legs, wings, and female ovipositor (3, 10, 34, 35). There are 5 types of labellar GRNs, defined by receptor expression, morphology, and functional properties: “water” neurons express pickpocket 28 (Ppk28) and encode low osmolarity; “sweet” neurons express gustatory receptor 64f (Gr64f) and respond to sugars and attractive (low) concentrations of sodium; “bitter” neurons express Gr66a and encode aversive compounds, including alkaloids and aversive concentrations of monovalent salts; “Ppk23^glut^” neurons are glutamatergic and respond to aversive concentrations of salts; and “IR94e” neurons express ionotropic receptor 94e (IR94e) and respond to salts and some amino acids (36–44). Labellar GRNs converge on a brain region called the subesophageal zone (SEZ), where a dense network of local neurons performs functions including sensory integration, gain control, and modulation (3, 45–51). GRNs also synapse with various classes of projection neurons that have been denoted as taste projection neurons (TPNs) or gustatory projection neurons (GPNs) in the SEZ, and these neurons carry information to various brain regions like antennal mechanosensory and motor center (AMMC), MB, and LH (3, 52–55). The distributed nature of taste processing is highlighted by a recent study demonstrating the divergence of sweet taste information in second-order neurons promoting different types of behaviors (56).

Despite the identification of various individual second-order *Drosophila* taste neurons and the lower dimensionality of taste identity coding relative to olfaction, a global organizing framework on which to analyze early taste processing principles remains elusive. With guidance from fly connectome data, we adopted a general approach combining connectomic analyses and functional imaging to investigate early taste organization principles across different taste qualities (57–60). We focused on taste coding and connectivity between three neuron types: GRNs (sensory neurons), LNs (interneurons), and TPNs (projection neurons). We uncover diverse tuning and connectivity of both GABAergic and cholinergic LNs. Moreover, we find that many TPNs are third order, indirectly receiving taste inputs through a layer of cholinergic LNs. We present evidence that cholinergic LNs may serve as nodes to integrate signals, including satiety information, and relay processed information to a set of morphologically similar TPNs. Thus, taste LNs may provide global feedback signals, as in olfaction, and also serve as key organizational units within fly taste circuits.

## Results

### Putative TPNs receive largely distinct inputs from each type of GRNs

To investigate the relationship between GRNs and their postsynaptic TPNs, we began by identifying GRNs and putative TPNs from the Flywire whole fly brain connectome dataset (57, 58, 61). GRNs in both hemispheres were identified and classified based on previously characterized morphology, recent GRN characterization by Engert et al. (2022), predicted neurotransmitter expression (62), and annotations submitted by community users (Figure S1A, see Methods for details) (40, 50, 51, 57–59, 62–64). We chose to restrict GRN divisions to 4 classes that we could separate with high confidence. “Bitter” and “IR94e” projections have unequivocally unique morphology, which was used to select these classes. Although, “sweet”, “water”, and “Ppk23^glut^” neurons all have similar morphologies, Ppk23^glut^ neurons were separable based on their predicted glutamatergic identity. This left a final “sweet/water” class, which we conservatively left as one group because we could not confidently separate sweet and water projections.

Putative TPNs were initially identified in Flywire based on two criteria: being directly postsynaptic to at least one GRN, and projecting to the Superior Medial Protocerebrum (SMP), the Superior Lateral Protocerebrum (SLP), or the lateral horn (LH). We restricted our TPN set to neurons projecting to these areas for the sake of relative simplicity; however, we acknowledge that putative TPNs projecting to other areas, such as the AMMC, also exist (55). The group of 57 identified TPNs will be referred to as “directly connected” TPNs going forward, to distinguish them from “indirectly connected” TPNs discussed later.

We next performed hierarchical clustering of GRNs based on their connectivity with directly connected TPNs (See Methods for details and Figure S1C for distance matrix). 10 clusters were chosen based on maximum separation indicated by the Silhouette score (Figure S1B). Within most of the clusters, GRN identities are relatively homogeneous. 6 clusters contain a single GRN identity, 3 clusters contain mixtures of two GRN identities but are primarily dominated by one, and 1 cluster contains a mixture of three GRN identities. The clustering result implies that different functional classes of GRNs differed in their downstream connectivity with putative TPNs (Figure 1A, B, and Figure S1D). To validate, we measured the pairwise cosine similarity of each pair of GRNs with respect to their connectivity with directly connected TPNs. As expected, this revealed higher similarity scores within each GRN class than between classes (Figure 1C). Replotting this data as the cosine similarity scores within each GRN class and between each pair of classes revealed heterogeneity of downstream connectivity within a GRN class (Figure 1D), and a distribution more centered on 0 for between class comparisons (Figure 1E). This further supports the notion that different classes of GRNs show largely distinctive downstream TPN targets.

**Figure 1.**
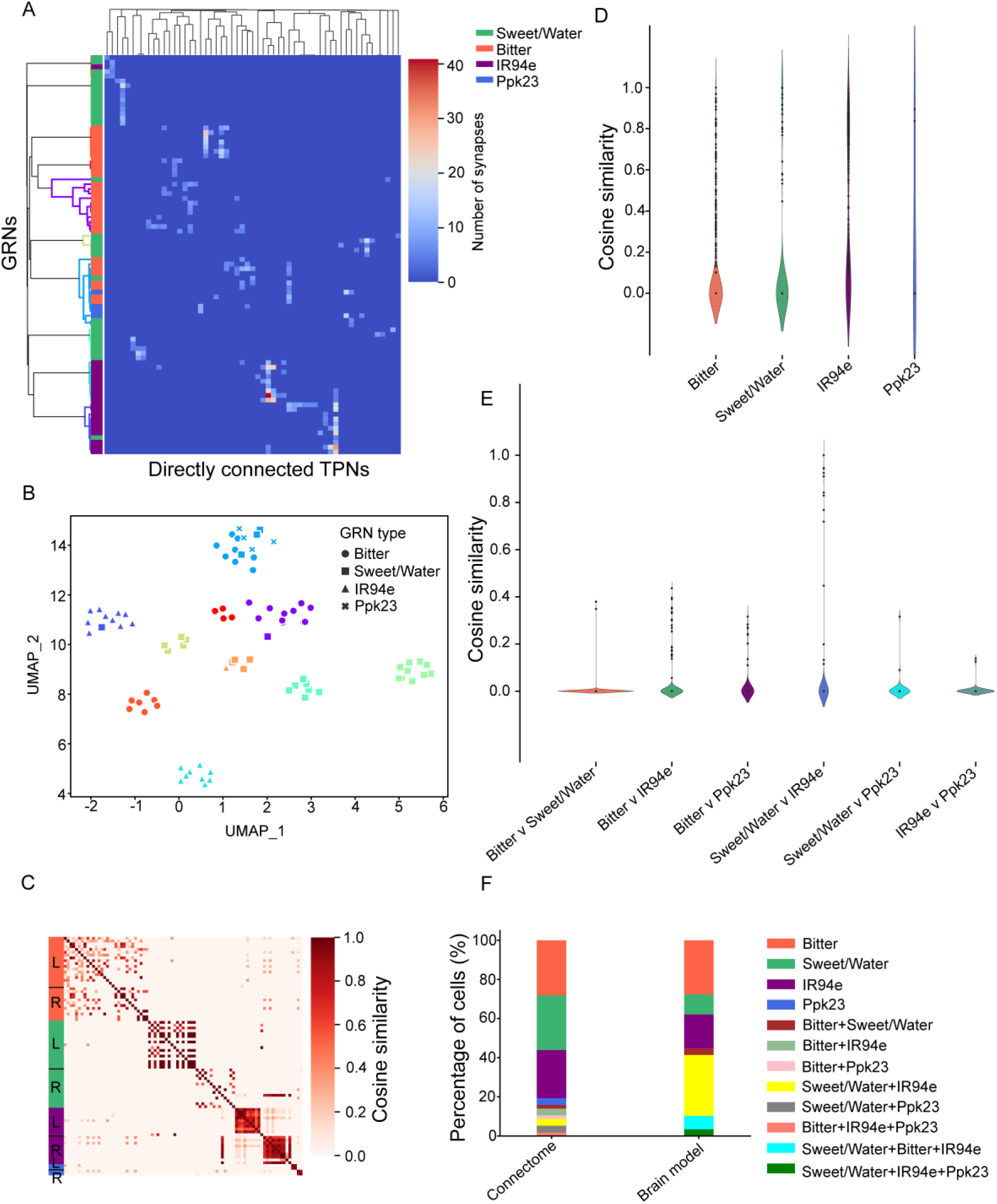
TPNs receive segregated inputs from each GRN type. **A)** Clustermap from hierarchical clustering demonstrating the raw connectivity between GRNs and putative TPNs. Dendrogram on the y-axis is color-coded based on the cluster labels in (B). **B)** UMAP embedding of clustered GRNs based on their putative TPN connectivity. **C)** Pairwise cosine similarity matrix of GRNs organized by their identity. Orange represents bitter GRNs, green represents sweet/water GRNs, purple represents IR94e GRNs, and blue represents Ppk23^glut^ GRNs. The color coding will be consistent in the rest of the figures. L = left side of the brain; R = right side of the brain. **D-E)** Pairwise cosine similarity within and across GRN types. The Kruskal Wallis test with Dunn’s posthoc test were performed and no statistical significance was found between any comparison. **F)** Comparison of cell proportions that receive different types of GRN inputs between two different measures: the connectome that predicts input based on direct connectivity and the brain model.

We noticed that many putative TPNs were connected weakly to the GRNs in the connectome, raising the possibility that some of them may not be taste responsive (Figure 1A). Moreover, the tuning of each TPN could also be influenced by non-receptor neurons that are downstream of the GRNs. We used a recently developed leaky integrate-and-fire model by Shiu and colleagues, hereon referred to as the “brain model”, to predict the activity of the putative directly connected TPNs (60). We initially ran the model by assessing GRNs of each type separately as input neurons and activating those neurons at 10-200 Hz with increments of 10 Hz (60). 42 out of 57 putative TPNs in our collection responded to some level of GRN stimulation (Figure S1E). Because the range of stimulation frequencies goes beyond what has been observed empirically via GRN recordings, we then restricted the stimulation range by setting the maximum activation level of each GRN class based on the plateau point where the number of neurons activated no longer increases rapidly (Figure S1F). 80 Hz was selected for sweet, water, and bitter GRNs activation experiments, which aligns well with previous electrophysiology recordings of 100 mM sucrose and 1 mM lobeline stimulation (64, 65). In the absence of much data on the physiological range IR94e GRN activation, we selected 150 Hz based on the rise in downstream activity up to this point. The entire 200 Hz range was included for Ppk23^glut^ GRNs because glutamate was predicted to be inhibitory in the model, resulting in only rare activation of downstream targets. Based on these modified stimulation parameters, 29 TPNs showed predicted taste-evoked activity and were analyzed further. We then compared the brain model predictions of tuning for this group of TPNs to the tuning predicted solely by annotating the class(es) of presynaptic GRNs. Compared to the tuning predicted by directly connected GRN type(s) in the connectome, the model predicted a higher proportion of the TPNs to be multiply tuned (Figure 1F).

One possible explanation for the broader TPN tuning predicted by the model is that taste-responsive local SEZ interneurons also provide excitatory drive to TPNs, thereby comprising an additional “indirect” link between GRN and TPN activation. To broaden our sampling of TPNs to include those indirectly connected via interneurons, we examined all the cholinergic LNs that are directly postsynaptic to GRNs, and collected putative TPNs that were postsynaptic to these interneurons. In all, we identified 155 indirectly connected putative TPNs, 23 of which also received direct inputs from the GRNs (Figure S2A). Based on the brain model, 38 out of the 155 putative TPNs showed taste responses (Figure S2B, S2C, S2D). 15 out of the 38 responsive TPNs also received direct inputs from GRNs, and were part of the directly connected TPN set analyzed above. Thus, 23 putative TPNs were predicted to rely completely on interneurons for their activation. This modeling data suggests an organization to the taste system that differs from the *Drosophila* olfactory system, where PNs generally all receive direct input from ORNs in the antennal lobe (20, 21). We hypothesize that the *Drosophila* local taste interneurons in the SEZ perform taste integration and also serve as nodes to distribute taste information in different circuits.

### Directly and indirectly connected TPNs selected from the connectome are activated by tastants

Due to the apparent complexity of the *Drosophila* taste system, we sought to verify the taste responses of both directly and indirectly connected putative TPNs. We selected three TPN cell types based on tuning predicted by the connectome and the availability of driver lines. We then used GCaMP imaging to examine their responses to different tastes.

As an example of a primarily indirectly connected TPN, we selected “peafowl” a neuron type identified in a collection of previously published SEZ split-Gal4 drivers (66). From the connectome, the right stereopair of peafowl receives 7 synaptic inputs from one sweet/water GRN, and the left stereopair of peafowl does not connect directly to any labellar GRNs. However, the neurons were predicted by the brain model to be activated by sweet/water GRNs, as well as IR94e GRNs stimulated at high frequency. Using a panel of tastants representing different modalities, we found that peafowl neurons respond primarily to sucrose, but the responses are variable across subjects (Figure 2A and 2B). Peafowl neurons also show variable responses to other types of sweet sugars such as glucose and arabinose (Figure 2C). Moreover, the responses to sucrose are dose dependent (Figure 2D; One-way repeated measure ANOVA p<0.05).

**Figure 2.**
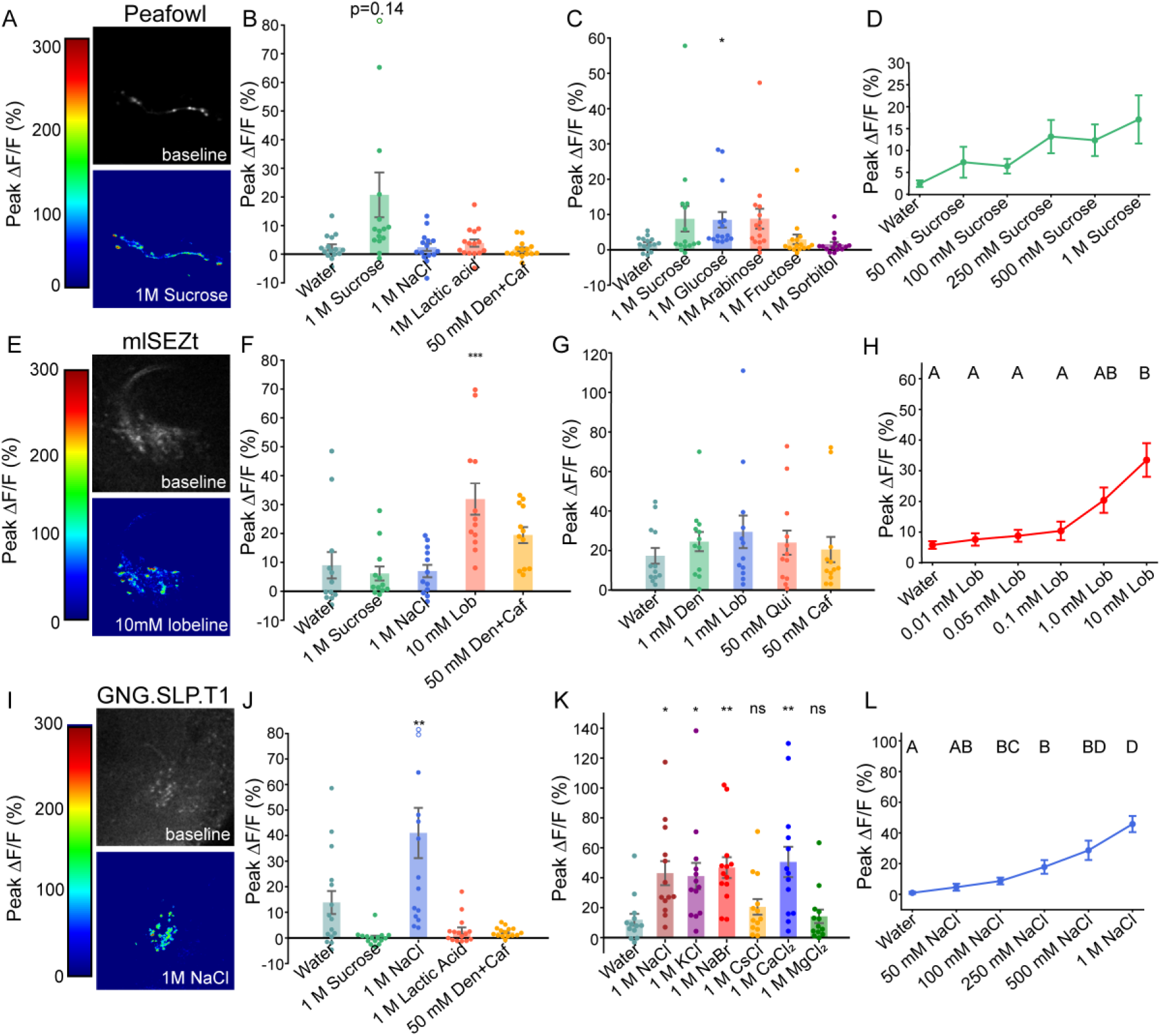
Example TPNs are taste responsive. **A)** Peak ΔF/F of peafowl projections in response to 1 M sucrose compared to baseline. **B-D)** Peak ΔF/F of peafowl projections in response to a panel of tastants with different taste modalities (B), different sugars (C), and increasing doses of sucrose (D), respectively. **E)** Peak ΔF/F of mlSEZt projections in response to 10 mM lobeline compared to baseline. **F-H)** Peak ΔF/F of mlSEZt neurons in response to different taste modalities (F), different bitter tastants (G), and increasing doses of lobeline (H), respectively. **I)** Peak ΔF/F of GNG.SLP.T1 projections in response to 1 M NaCl compared to baseline. **J-L)** Peak ΔF/F of GNG.SLP.T1 projections in response to different taste modalities (J), different salts (K), and increasing doses of NaCl, respectively (L), respectively. One-way ANOVA and Dunnet’s posthoc test compared to water were performed in B, C, F, G, J, K. One-way ANOVA with Tukey’s multiple comparison were performed in D, H, L. N=12-15 replicates, *p < 0.05, **p < 0.01, ***p < 0.001.

The second TPN we examined is predicted to be the same as previously identified mlSEZt projection neurons due to their morphological similarity and bitter-specific tuning (Figure 2E, 2F, and Figure S3E-F; 53, 66). According to the connectome, some neurons in this population, labeled by the split-Gal4 driver R43D01-p65; R29F12-DBD, are second order neurons that form direct synapses with bitter GRNs, while others are indirectly connected via cholinergic local interneurons. Consistent with previous reports, we found that the mlSEZt neurons respond to multiple bitter tastants, exhibiting clear peaks corresponding to the onset and removal of the stimulus (Figure 2G and Figure S3G; 67). The response to lobeline was also dose dependent (Figure 2H and Figure S3H; One-way repeated measure ANOVA p<0.05).

The third TPN we tested was GNG.SLP.T1, which receives inputs from IR94e GRNs (42). We selected GNG.SLP.T1 neurons because they are an example of cells that receive almost entirely direct inputs from GRNs. Among the panel of tastants examined, the GNG.SLP.T1 neurons responded primarily to salts and had a minor response to water (Figure 2I-K and Figure S3I-K). We observed responses to both monovalent and divalent salts, and dose-dependent activation by NaCl (Figure 2K, 2L, and Figure S3K-L). Although IR94e neurons are known to have some salt sensitivity, salts also activate other GRN types, such as those labelled by Gr64f, Gr66a, and Ppk23^glut^ (40–42). To rule out the possibility that GNG.SLP.T1 receives input from non-IR94e GRNs, we first confirmed that salt responses depend on IR76b, one of two co-receptors required for salt responses in flies, demonstrating that GNG.SLP.T1 salt activation is a genuine taste response (40, 41, 43, 69, 70) (Figure S4A and S4B). Based on the lack of sweet and bitter responses in the GNG.SLP.T1 neurons, we inferred that inputs are unlikely from Gr64f (sweet) or Gr66a (bitter) GRNs. Moreover, mutants for *IR7c*, which is required for monovalent salt responses in Ppk23^glut^ GRNs (41), showed normal salt-evoked activity in GNG.SLP.T1 (Figure S4C and S4D). *IR94e* mutants also showed normal GNG.SLP.T1 activity (Figure S4E and S4F); however, imaging of IR94e sensory neurons revealed intact salt responses in *IR94e* mutants, suggesting the existence of additional salt-sensing IRs in this population (Figure S4G and S4H). Overall, our imaging of three example TPN types reveals that in all three cases, TPN tuning mirrors the tuning of the GRNs either directly presynaptic or indirectly connected via one cholinergic interneuron.

### LNs integrate sensory information across multiple GRN types

To investigate the possible roles of LNs in the transfer of taste information from GRNs to TPNs, we began by performing hierarchical clustering of GRNs based on their postsynaptic connectivity with either cholinergic or GABAergic LNs (See Methods for details and Figure S5B and S6B for distance matrices). The raw number of synapses between each GRN and each of 82 cholinergic LNs illustrates a diversity of LNs based on their patterns of GRN inputs (Figure 3A). However, clustering of GRNs produced separation of sweet/water and bitter GRNs with some intermixing of Ppk23^glut^ and IR94e types (Figure 3A). This view is supported by UMAP embedding of the GRNs, where the maximized silhouette score resulted in 11 clusters that were mostly homogeneous for GRN class (Figure 3B, Figure S5A-C). One exception is Ppk23^glut^ GRNs, which appear in many clusters, suggesting that they may share common downstream circuits with other types of GRNs (Figure S5C, blue bars). Examining pairwise cosine similarity between GRNs also supports the idea that GRN connectivity with cholinergic LNs is more similar within a GRN class than between classes, and that some Ppk23^glut^ GRNs show connectivity similar to either sweet/water or bitter classes (Figure 3C-E). However, pairwise cosine similarity within GRN and between GRN classes shows substantial heterogeneity, suggesting the existence of many different sets of inputs to cholinergic LNs. Notably, IR94e GRNs show generally low cosine similarity with other GRNs, likely because they are only sparsely connected to cholinergic LNs (Figure 3A), instead forming direct synapses with TPNs (Figure 1A).

**Figure 3:**
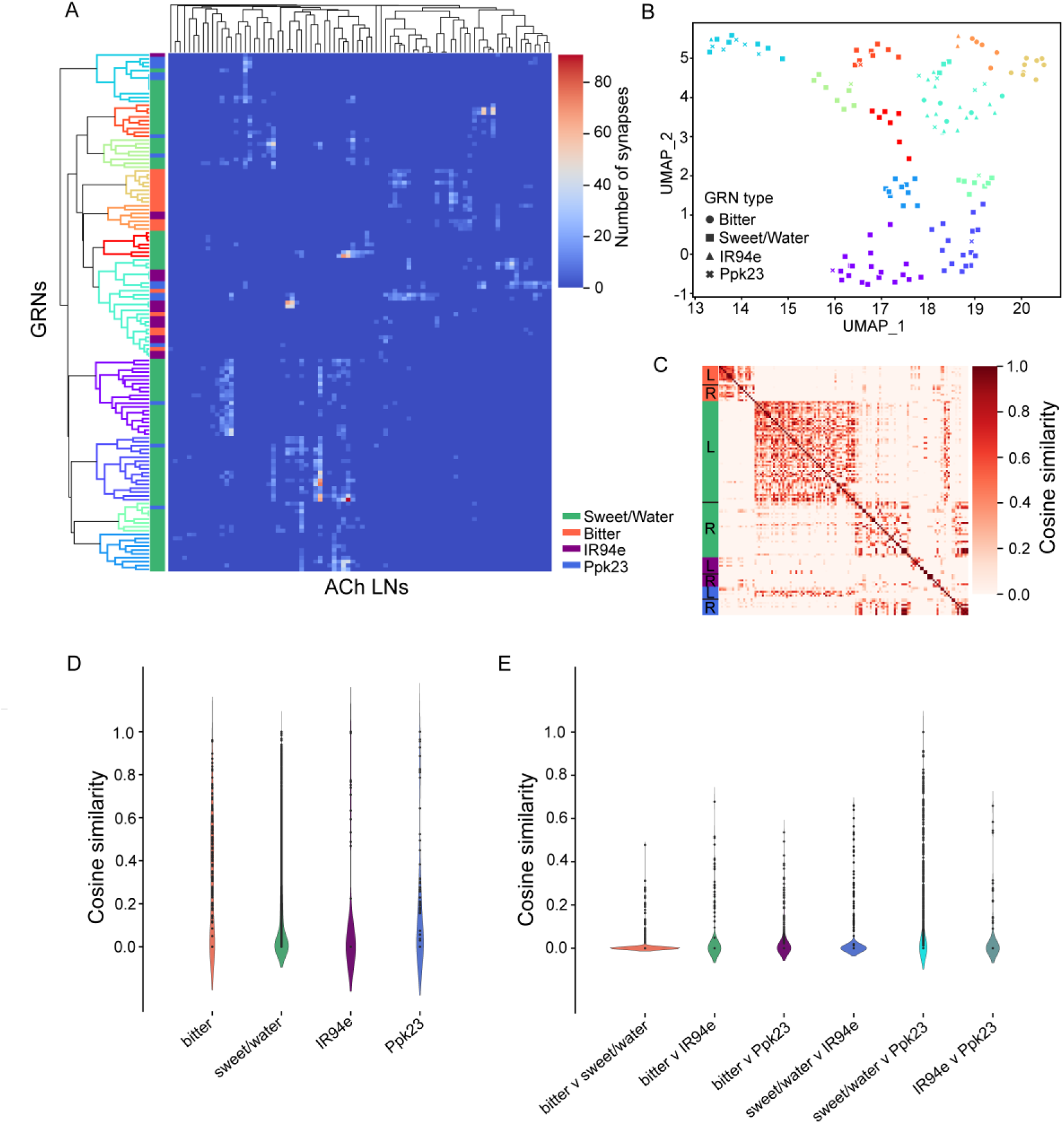
Connectomics analyses show integration of GRN inputs by cholinergic LNs. **A)** Clustermap from hierarchical clustering showing raw connectivity between GRNs and their downstream cholinergic local interneurons. Dendrogram on the y-axis is color-coded based on the cluster labels in B). **B)** UMAP embedding of GRN clusters based on their downstream cholinergic LNs connectivity. **C)** Pairwise cosine similarity between GRNs based on their downstream cholinergic LNs connectivity. **D-E)** Pairwise cosine similarity within and across types of GRNs, respectively. Kruskal Wallis with Dunn’s posthoc test were performed and no statistical significance was found between any comparison.

Similar to cholinergic LNs, the raw number of synaptic inputs from GRNs to the 50 identified GABAergic LNs suggests general segregation between sweet/water and bitter GRN classes (Figure 4A). However, we also observed strikingly similar connectivity between bitter and IR94e GRNs, which are both heavily connected to two specific GABAergic LNs. This observation aligns with previous studies suggesting that IR94e and bitter GRNs inhibit feeding motor programs via GABAergic LNs (42, 60). UMAP embedding revealed 8 clusters of GRNs with a maximized silhouette score (Figure 4B and Figure S6A). This analysis also placed bitter and IR94e GRNs within the same clusters, driving the overall separation of GRN types to be lower than what we observed for cholinergic LNs (Figure S5C, S6C). Additionally, sweet/water and Ppk23^glut^ GRNs often fall under the same clusters, similar to our clustering of GRNs based on cholinergic LN connectivity (Figure 3). As expected, bitter and IR94e, and sweet/water and Ppk23^glut^ GRNs share high pairwise cosine similarity of downstream GABAergic LN connectivity (Figure 4C and 4E). This is particularly strong for IR94e GRNs, which have high levels of homogeneity based on their connection to the two GABAergic interneurons shared with bitter inputs (Figure 4D and 4A).

**Figure 4:**
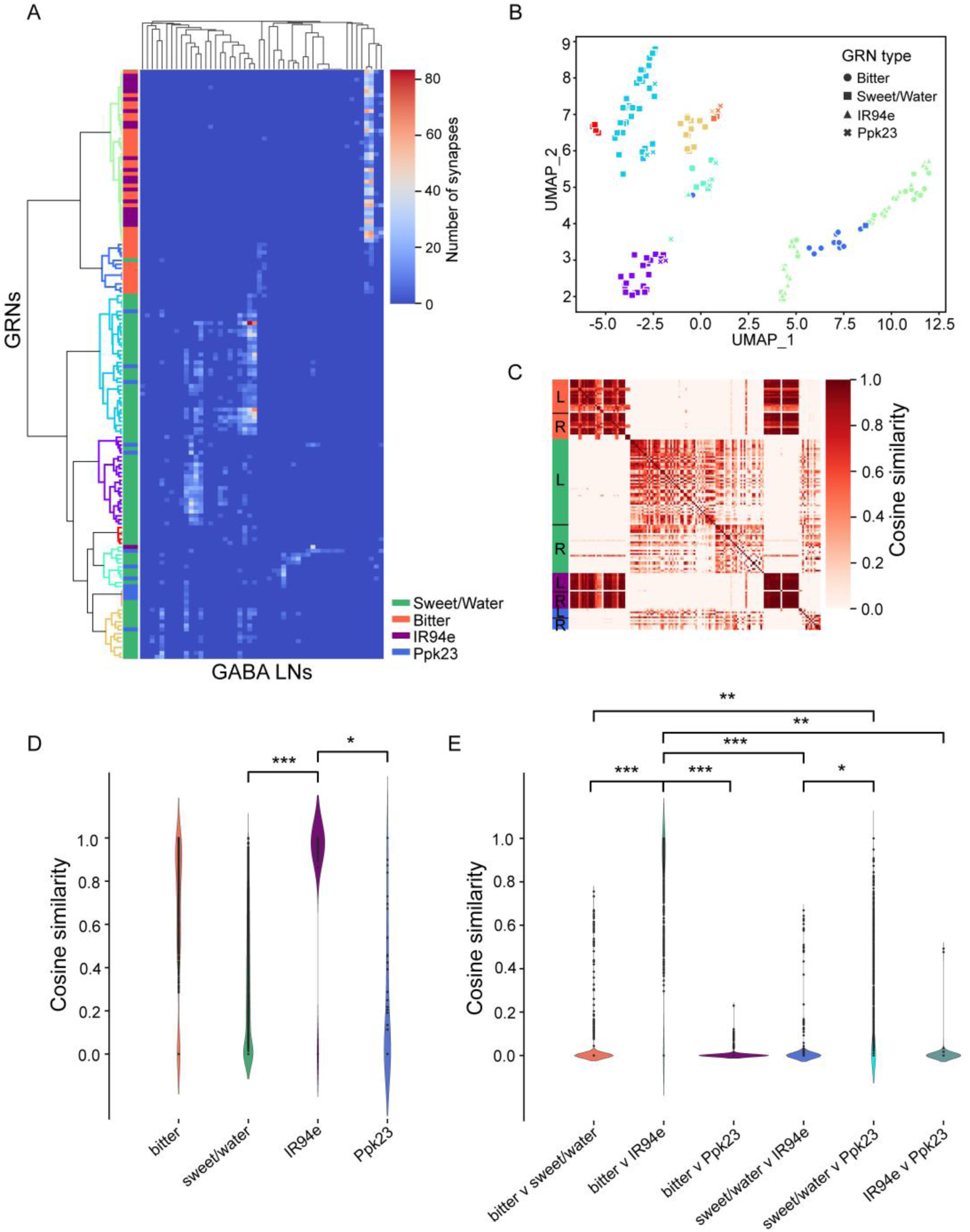
Connectomics analyses show integration of GRN inputs by GABAergic LNs. **A)** Clustermap from hierarchical clustering showing raw connectivity between GRNs and their downstream GABAergic LNs. Dendrogram on the y-axis is color-coded based on the cluster labels in B). **B)** UMAP embedding of GRN clusters based on their downstream GABAergic LNs connectivity. **C)** Pairwise cosine similarity between GRNs based on their downstream GABAergic LNs connectivity. **D-E)** Pairwise cosine similarity within and across types of GRNs, respectively. Kruskal Wallis with Dunn’s posthoc test were performed. *p < 0.05, **p < 0.01, ***p < 0.001.

### Calcium imaging of SEZ cell bodies shows mixture of singly and multiply tuned cells

Our analysis of GRN-to-LN connectivity suggests that many LNs receive inputs from a single GRN type, while some likely integrate inputs across multiple modalities. To test this prediction empirically, we performed calcium imaging of cholinergic or GABAergic cell bodies in the SEZ by driving GCaMP6s under the control of Cha-Gal4 or GAD1-Gal4, respectively. Four different natural taste stimuli were chosen to activate specific GRN types based on previous studies: water to activate Ppk28 GRNs; 1 M sucrose to activate Gr64f GRNs; 50 mM CaCl_2_ to activate Ppk23^glut^ GRNs; and a mixture of 50 mM denatonium and 50 mM caffeine to activate bitter GRNs across different sensilla (68, 71). To specifically activate IR94e neurons, we expressed exogenous P2X_2_ receptors in the IR94e GRNs and stimulated with 100 mM ATP.

For each genotype, the tuning profiles of ∼600 individual cell bodies, measured across 6 individual flies, were analyzed after applying a 10% threshold criterion for considering a cell to have responded to a given stimulus (Figure 5A and 5E). For cholinergic neurons, 62% of cells were singly tuned, while 38% were multiply tuned (Figure 5A and 5B). The doubly tuned cells included most of the possible taste combinations (Figure 5C). As expected, there was only a small proportion of cells responding to both 1 M sucrose and the bitter mixture, suggesting that appetitive and aversive pathways are segregated at this level (Figure 5A and 5C). Among GABAergic cells, 67% were singly tuned and 33% were multiply tuned (Figure 5E and 5F). Again, the multiply tuned cells had a variety of tunings, and the proportion of cells that responded to both 1 M sucrose and bitter mixture was low (Figure 5E and 5G).

**Figure 5:**
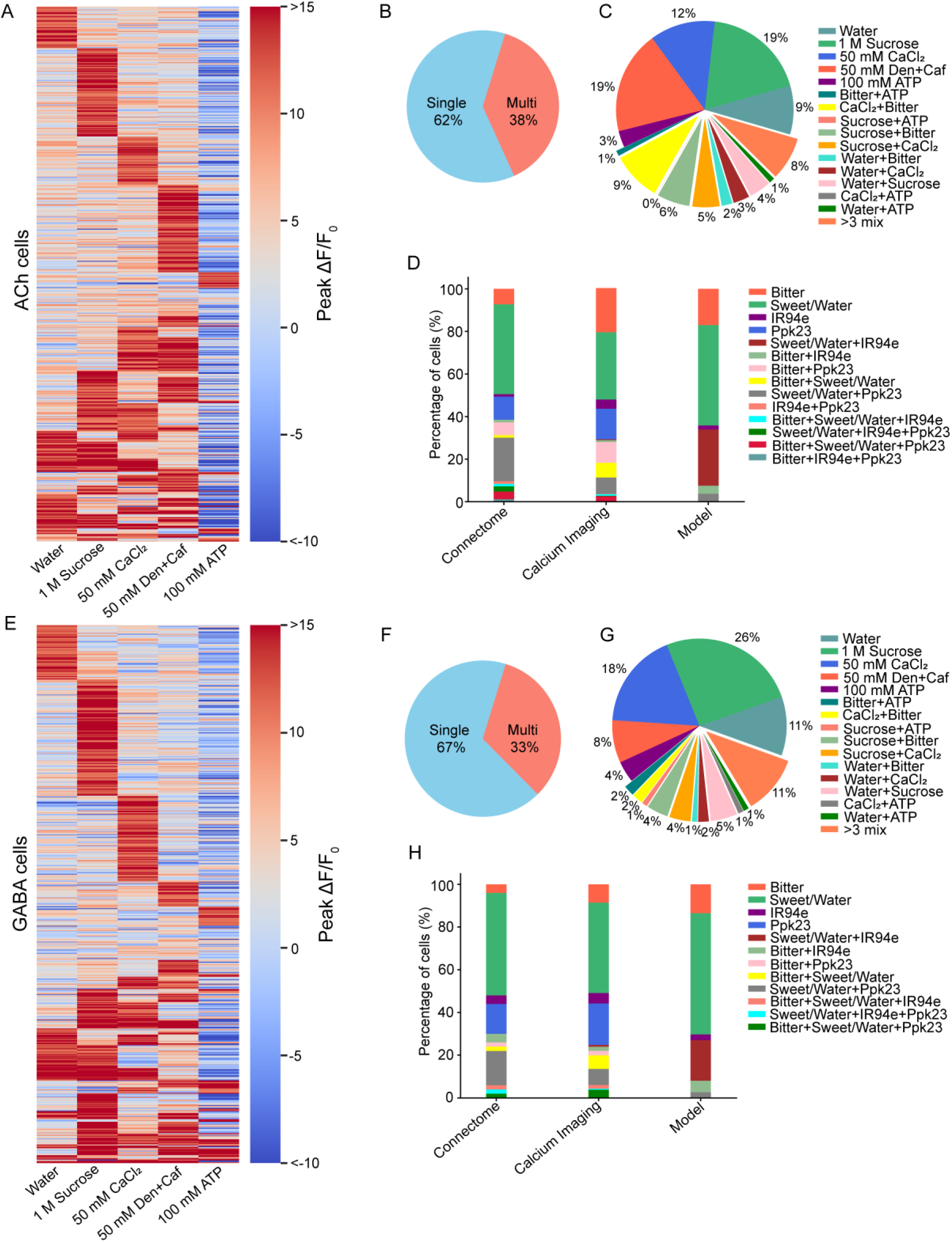
Calcium imaging of SEZ cholinergic and GABAergic cells supports taste sensory integration. **A)** Heatmap of cholinergic cells that respond to any tastants with 10% or higher peak ΔF/F. **B)** Proportion of singly vs multiply tuned cholinergic cells. **C)** Detailed proportion of singly and doubly tuned cholinergic cells with different response profiles. **D)** Comparison of cholinergic cell proportions that receive different types of GRN inputs with three different measures: inputs based on direct connectivity with the GRNs in the connectome, inputs based on calcium imaging response profiles, and the GRN activation in the brain model. **E)** Heatmap of GABAergic cells that respond to any tastants with 10% and higher Peak ΔF/F. **F)** Proportion of singly vs multiply tuned GABAergic cells. **G)** Detailed proportion of singly and doubly tuned GABAergic cells with different response profiles. **H)** Comparison of GABAergic cell proportions that receive different types of GRN inputs with three different measures as described in panel (D).

We next wondered how our calcium imaging results compared to predictions based on the connectome. Input GRN types were inferred for each neuron in our calcium imaging datasets and then compared to data from the connectome evaluated in two ways: classifying each LN’s direct input GRNs (“connectome”), and the GRN type(s) sufficient to activate each LN using the whole brain model (“model”). For cholinergic interneurons, the proportion of cells with each tuning type is similar among the three types of measurements, although the brain model predicts fewer LNs activated by Ppk23^glut^ GRNs, since glutamate is classified as an inhibitory neurotransmitter in the model (Figure 5D). Cholinergic cells responding by Ppk23^glut^ and sweet/water GRNs are also over-represented in calcium imaging relative to the brain model, although both are smaller than the proportion of LNs receiving inputs from both Ppk23^glut^ and sweet/water GRNs (Figure 5D, Figure S7A and S7B). Interestingly, the brain model predicts a large proportion of cholinergic LNs responding to sweet/water and IR94e GRNs, whereas this class is virtually absent in the calcium imaging data and connectome predictions (Figure 5D).

Like cholinergic cells, GABAergic LNs showed a strong consensus between the calcium imaging results and tuning predicted by direct GRN inputs or the brain model, with exceptions for the model’s low prediction of responses to Ppk23^glut^ and high prediction of mixed tuning between sweet/water and IR94e (Figure 5H, Figure S7C and S7D). Overall, our analyses suggest that the tuning of LNs is largely predictable by their direct inputs from GRNs. Moreover, consistent with a previous study by Harris et al. (48), most LNs are singly tuned, a minority respond to inputs from multiple GRN types, and cells responding to sweet and bitter are rare. The picture that emerges is one where LNs mostly provide feed-forward or feedback activity within a modality but LNs likely also contribute to integration between modalities.

### Subsets of cholinergic LNs are differentially modulated by internal state

GABAergic LNs in the fly olfactory system have well documented roles in presynaptic and postsynaptic gain control, which can operate within a glomerulus or between glomeruli (25, 26, 28, 29). Similar roles are likely in the taste system, where feedback inhibition could perform divisive normalization and mixture suppression between modalities (47). However, the putative role of cholinergic LNs in the taste system is less clear. One possibility is that they may serve as a site of modulation to integrate sensory signals with information about internal state. For example, IN1 neurons receive inputs from pharyngeal sweet neurons and their activity is modulated by hunger (45). Interestingly, hunger modulation is also selective to a subset of second-order neurons in the proboscis extension motor circuit (46). To investigate the potential role of labellar cholinergic LNs in taste modulation, we performed calcium imaging on two types of LNs driven by specific split-Gal4s: “Rattle” (66) and a set of neurons labeled by R18H03-AD; VT046048-DBD, which we believe to correspond to neurons previously named “marge” (Figure 6A and 6C).

**Figure 6:**
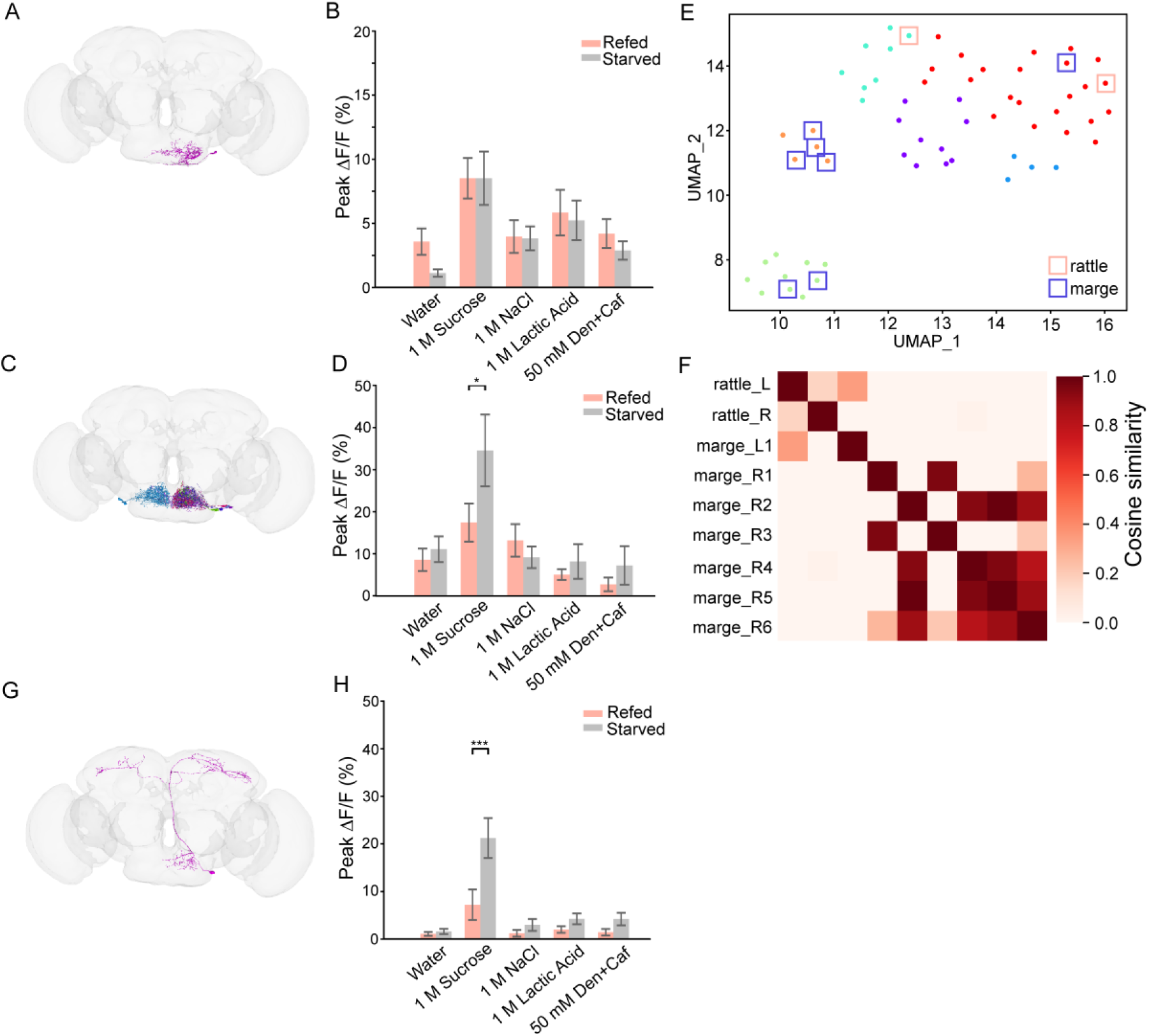
Cholinergic LNs show different responses to hunger modulation. **A)** Neuron skeleton of rattle LNs in the connectome. **B)** Calcium imaging of rattle with peak ΔF/F responses to tastants of different modalities in refed vs starved conditions. **C)** Connectome image of putative marge LNs. **D)** Calcium imaging of marge peak ΔF/F responses to tastants of different modalities in refed vs starved conditions. **E)** UMAP embedding of second-order cholinergic interneurons clusters with rattle and marge labelled. **F)** Pairwise cosine similarity matrix of rattle and marge based on the Gr64f/Ppk28 inputs. **G)** Connectome image of peafowl TPNs. H) Calcium imaging of peafowl peak ΔF/F responses to tastants of different modalities in refed vs starved conditions. Two-way ANOVA with Šídák’s multiple comparisons test were performed. N=15-18 flies, *p < 0.05, ***p< 0.001.

Calcium responses of the two neurons were measured following 24 h starvation with or without 1 h of refeeding with standard cornmeal medium just prior to imaging. While both neuron types responded to 1 M sucrose, only the response of marge neurons increased with starvation, relative to refed controls (Figure 6B and 6D, Figure S8A and S8B). Thus, rattle and marge LNs may represent organizational nodes to circuits with differential requirements for hunger modulation. To explore the source of this difference in modulation, we examined whether it could be derived from differential input from sweet/water GRNs. Hierarchical clustering was conducted on the cholinergic LNs based on their connectivity to sweet/water GRNs, and 6 clusters were selected based on the silhouette score (Figure 6E, Figure S8C and S8D). We identified one rattle LN on each side of the brain, while seven LNs were identified with marge-like morphology (1 on the left side and 6 on the right side). Six of the seven marge LNs cluster away from rattle LNs, suggesting that they have different sweet/water GRN connectivity (Figure 6E). Pairwise cosine similarity confirms that rattle and marge neurons receive different sweet/water GRN inputs, except for the one marge neuron in the left hemisphere (Figure 6F). Thus, it is possible that different sweet/water GRNs exhibit different levels of hunger modulation, and these differences are selectively passed to taste LNs. It is also possible that LNs represent the direct site of modulation from other hunger-sensitive pathways.

If each cholinergic LN serves as a node that organizes information to be relayed to projection neurons, we would expect hunger modulation seen in LNs to be passed on to downstream TPNs. Peafowl, a sweet-responsive TPN that receives negligible inputs directly from labellar sweet/water neurons, was found to be postsynaptic to half of the putative marge cholinergic interneurons in the connectome (Figure 6G). As expected, peafowl neuron activity is also modulated by hunger states, suggesting that modulation occurring at the level of, or upstream of, LNs is faithfully passed to TPNs (Figure 6H and Figure S8E).

### Indirectly connected TPNs show morphological organization

If cholinergic LNs serve as nodes where taste inputs are transmitted to distinct circuits, one prediction is that TPNs connecting to a particular cholinergic LN will be similar to each other. We performed hierarchical clustering on both axes of a connectivity matrix between cholinergic LNs and indirectly connected TPNs, with cholinergic LNs color-coded based on their inferred GRN inputs. This clustering shows that each cholinergic LN generally connects to a sparse set of downstream TPNs (Figure 7A, Figure S9A). Visual examination demonstrates apparent morphological similarity between the TPNs connecting to an individual cholinergic LN, suggesting that each cholinergic LN might be dedicated to propagating information to specific higher order circuits (Figure 7B). To quantitatively assess the organization, we used NBLAST to generate a similarity score for each pair of TPN skeletons postsynaptic to the same cholinergic LN. We then compared the average pairwise similarity score against 10 shuffled datasets where TPN pairs were selected regardless of whether they connected to the same cholinergic LN (72). The unshuffled set has a significantly higher average NBLAST similarity score than any of the shuffled sets, suggesting that the morphological similarity between TPNs connected to a given cholinergic LN is non-random (Figure 7C).

**Figure 7:**
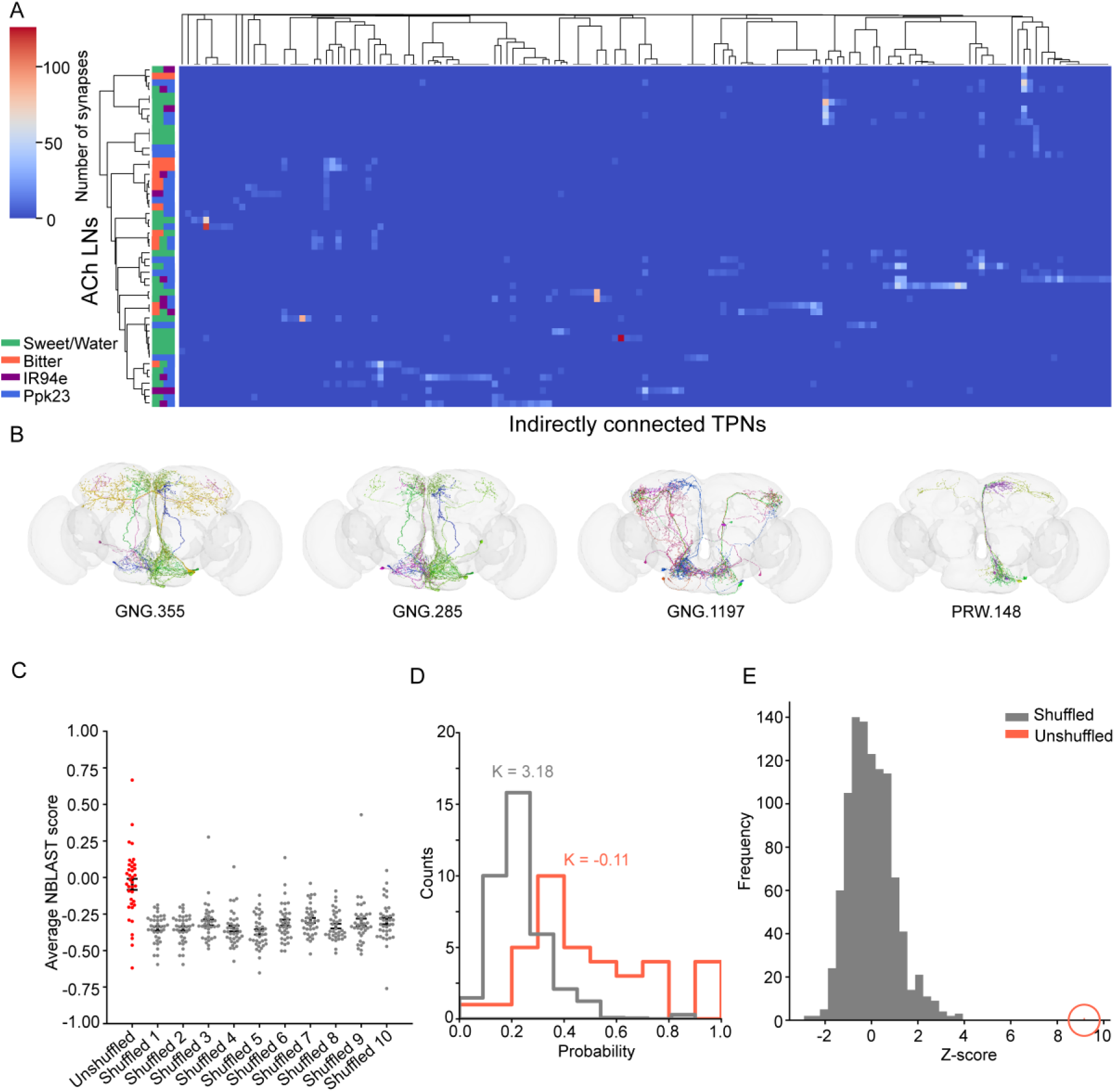
Third-order TPNs are organized based on morphological similarity. **A)** Clustermap showing raw connectivity between cholinergic LNs and third-order TPNs. The inferred GRN inputs for each cholinergic LN were color-coded on the y-axis. **B)** Connectome image of all TPNs postsynaptic to each example cholinergic LN. The TPN skeletons are postsynaptic to the cholinergic LNs indicated below each brain image. **C)** Average NBLAST scores of the collective TPNs postsynaptic to each cholinergic LN in unshuffled and 10 shuffled sets. Each data point represents the average NBLAST score of the TPNs postsynaptic to a given cholinergic LN. Student’s t-test was used to compare between unshuffled and each shuffled set, and significant differences of p<0.001 was found in all comparisons. **D)** Probability distribution of two TPNs belonging to the same morphological cluster for unshuffled and 1000 shuffled sets. K represents kurtosis. The probability distribution of the unshuffled set has a heavier tail. **E)** Mean probability distribution for unshuffled vs 1000 shuffled sets. Circle marks the mean probability for the unshuffled set.

To examine this idea further, we performed hierarchical clustering of the indirectly connected TPNs based on morphological similarity scores, generating 4 clusters (Figure S9C). Calculating cosine similarity between every pair of TPNs based on their connectivity with presynaptic cholinergic LNs revealed higher scores within each morphological cluster of TPNs, suggesting some organization (Figure S9B). We then calculated a score for each cholinergic LN that represents the probability that any randomly selected pair of its postsynaptic TPNs is comprised of two TPNs belonging to the same morphological cluster (see Methods for details). Descriptive statistics were used to compare the unshuffled data with 1000 sets of shuffled data, demonstrating that the mode and mean probability in the shuffled data occur around the expected value of 0.25, while the mean probability is shifted higher in the unshuffled dataset (Figure 7D). The Z-scores for the means of the 1000 shuffled data was calculated and the Z-score of the mean of the unshuffled data falls outside the distribution (Figure 7E). These results indicate that TPNs postsynaptic to a given cholinergic LN are morphologically more similar to each other than to the TPN population as a whole, demonstrating some degree of logical organization.

## Discussion

In this exploratory study, we examined the organizing principles of fly taste circuits at early steps of processing. Based on analyzing data from a whole brain connectome, we found third-order TPNs that do not directly form connections with the labellar GRNs, instead receiving inputs from local interneurons. This type of organization may facilitate integration of taste inputs with other modalities and with interoceptive signals. Moreover, each cholinergic LN is connected to morphologically similar TPNs, suggesting that cholinergic LNs function as hubs for information flowing to specific projection neuron target areas.

### Organization of different layers of taste neurons

One challenge we faced was the unequivocal categorization of GRNs into known functional classes. Bitter and Ir94e GRNs were easy to identify based on their unique morphological features, but the morphology of sweet, water, and Ppk23^glut^ GRNs are all very similar. We chose to separate Ppk23^glut^ GRNs based on neurotransmitter prediction. However, these predictions remain imperfect, leading to the likelihood that some GRNs are misclassified. We also chose to leave sweet/water GRNs as one class, an approach that differs from two previous studies. Engert et al. separated sweet (Gr64f) and water (Ppk28) GRNs using morphological and connectivity clustering (63), while a follow-up study by Shiu et al. (2023) assigned water GRNs based on connectivity with a second-order neuron (“usnea”) that is only water responsive (46, 60). Although both these approaches are reasonable, further studies exploring the morphological differences within a GRN class (for example, sweet neurons from different sensilla) and their connectivity will be needed to confirm the predictions. Nonetheless, while the purpose of these studies – making accurate predictions of information flow and behaviour based on the connectivity of neurons – requires the separation of different GRN types, it likely tolerates minor misidentification. Moreover, the observation that sweet and water inputs activate some shared downstream circuitry adds to the justification of our conservative approach of leaving sweet/water GRNs as a single class (36, 38, 46).

From the connectome, we found that many putative TPNs are third order, connecting indirectly with GRNs via a layer of cholinergic LNs. We confirmed that some of these putative TPNs are taste responsive using the brain model developed by Shiu et al. (2023) and using calcium imaging to look at the responses of peafowl, which primarily receives input from the cholinergic LNs rather than directly from GRNs. This organization differs from the olfactory system, in which all the PNs are directly postsynaptic to ORNs, with local excitatory and inhibitory neurons tuning inputs from the ORNs and outputs of the PNs (8, 20, 21). Perhaps this different organization reflects distinctions in the coding and outputs of taste versus smell. Taste coding is a much lower dimensional problem than olfactory coding, with pure tastants generally activating one of only a handful of functionally distinct sensory neuron classes (3, 4, 34). At the same time, taste circuits also appear more distributed across the fly brain, compared to olfactory projection neurons that have two primary targets in the Mushroom Bodies and Lateral Horn (8, 21). Thus, cholinergic LNs in the taste system may serve as individual nodes to organize streams of circuits that convey taste information to different parts of the fly brain. Our observation that that each cholinergic LN has above-chance likelihood of synapsing onto morphologically similar TPNs further supports this idea. The parallel streams of taste information early in the circuit can facilitate downstream decoding and help with the generation of appropriate behaviors due to less interference, while also imparting specific gain control mechanisms or modulatory controls over individual targets. Divergence of taste circuits is also supported by a recent behavioral study, in which a single taste modality drives behaviorally distinct responses (56).

In our NBLAST analysis, we noticed that many pairwise comparisons of TPN morphology generated negative scores even though we took the means of the forward and reverse comparison scores. This indicates that some pairs of TPNs have NBLAST similarity scores below the natural cut-off of the average pairwise comparison scores from random pairs of neurons in the brain (72). However, previous studies examining olfactory PN morphology all have above zero pairwise similarity score (72, 73). The source of this discrepancy is likely the spatially distributed nature of neuron backbones of the TPNs, whereas the backbones of many PNs are spatially restricted. Moreover, we found that stereopairs of TPNs have lower than expected pairwise similarity scores, even though they are mirror image of each other and possibly carry the same functional information. This limitation has potentially led to us underestimating the similarity scores of each TPN group during quantification. Future improvement on the algorithm considering the stereopairs is important for generating more accurate and meaningful neuron comparisons.

### Taste coding in second- and third-order neurons

Although the TPNs we examined with calcium imaging are all singly tuned to one taste modality, evidence from the connectome, the brain model, and a study of second-order TPN responses, suggest the presence of multiply tuned TPNs (74). However, the multiply tuned TPNs predicted from the connectome and brain model generally do not receive inputs from both sweet and bitter GRNs, demonstrating strong separation of these two modalities. The same can be said of taste LNs in the SEZ, which we were able to survey more broadly using calcium imaging. Again, most cholinergic and GABAergic LNs appear to be singly tuned, and few respond to both sweet and bitter taste. One limitation of this experiment is that some of the cell bodies we imaged could be TPNs with cell bodies residing in the SEZ, which may have skewed our measurements of the proportion of cells falling into each tuning group. Another limitation is that we applied tastants at concentrations above physiological relevance to parse the GRN inputs to LNs with cell bodies residing in the SEZ. While this strategy to maximize signal-to-noise has been standard practice in the field, it may also be expected to increase the number of multiply tuned cells, as has been observed in taste responses of mouse geniculate ganglion neurons (19). Lastly, the 10% ΔF/F threshold we applied for determining responsive cells is somewhat arbitrary, as some cells have peak ΔF/F responses slightly under 10%, rendering them unresponsive under our criterion. Therefore, this limitation can cause the tuning profiles to be different from the actual tuning of the cells, and the inferred GRN inputs to these cells can also be inaccurate. Despite the limitations, our results are consistent with the prior imaging studies suggesting that a subset of GABAergic LNs responds to both sweet and bitter stimuli, although these represent a small fraction of the taste-responsive SEZ cell population (47, 48).

Overall, our results support the idea that taste coding adheres to neither a strict labeled line nor a primarily combinatorial model. In the distributed taste circuits of the fly brain, breadth of tuning may reflect the particular function of specific circuits. For example, co-activation of many feeding circuit neurons by sugar and water may increase efficiency of coding stimuli that elicit consumption (46), while the same may not be true in circuits impacting other behaviors such as oviposition. It’s also important to note that we examined coding from the perspective of stimuli that activate single GRN types. Other stimuli that activate multiple GRN types, such as salt and acids, may be represented in a more complex way in the brain. Nonetheless, the strong separation of sweet from bitter is a theme that remains true across ours and several other studies (46, 48, 60), and likely reflects both the specificity of these stimuli for individual GRN types and their consistently opposing valence across different behaviors (3, 4, 34, 48).

### Local cholinergic interneurons as a site of modulation

The suggestion that each cholinergic LN serves as a node to convey information through distinct parallel streams led us to consider the functional relevance of this organization. We found two cholinergic local interneurons, rattle and marge, that are both postsynaptic to sweet GRNs but are differently modulated by hunger state. These two neurons also show distinct connectivity with sweet GRNs, suggesting that there are subdivisions within the sweet pathway as far out as the sensory neurons. Notably, although labellar sweet GRNs are known to be modulated by hunger states via dopamine signaling, whether this is true of all (or even most) sweet GRNs is still unknown (75–77). An alternative explanation is that interoceptive neurons receiving information about the nutrient levels may signal directly to specific LNs. Finally, it is possible that non-linear transformations at connections between sweet GRNs and different LNs may emphasize small differences in GRN firing in some LNs and de-emphasize them in others. Regardless of the mechanism of modulation, having different levels of modulation across parallel sweet circuits may enable maximum flexibility in some while maintaining robust assessment of taste identity and concentration in others.

### Materials and Methods Flies

Flies were reared on standard cornmeal medium and kept at 25 °C in 60% humidity under 12-hour light and dark cycles. See Table S1 for detailed genotype information.

### Calcium imaging

Mated females aged 3-10 days were used. Flies were fed normally before imaging unless otherwise noted. For refed and starved experiments, flies were split into two groups and starved on 1% agar-only medium for 24 hrs. One group was refed for an hour after starvation. *In vivo* calcium imaging under a 25X water-immersive objective using a Leica SP5 II Confocal microscope was performed as previously described (40, 41, 78). Flies were anesthetized, amputated, and mounted on a custom-made chamber. Nail polish was applied on the retina of the fly to secure the fly head and expose the dorsal brain region for TPN imaging. For imaging the SEZ, nail polish was applied on the dorsal side of the head. The fly proboscis was fixed in an extended position using wax. Mounted flies were dissected after > 30 min recovery. AHL with Mg^2+^ and Ca^2+^ ions was applied on the fly head during dissection (Jaeger et al., 2018). The anterior cuticle was removed and the fat bodies and air sacs were cleared from the imaging window.

For TPN and interneuron 2-dimensional (xy) time series acquisition, argon laser power was set to 6-10% with pinhole opened to 209.95 µm. Axonal projections were imaged with a window of 4-8X zoom and 256×256 resolution was used for acquisition. Acquisition line speed was set to 8000 Hz with line average accumulation set to 2-4. The frame rate was 36.4-69.2 ms/f. 10 s baseline was acquired before stimulation, and each stimulation lasts for 5 s.

For volumetric imaging of ACh and GABA neurons in the SEZ, argon laser power was set to 6% with pinhole opened to 55.86 µm. Cell bodies within the SEZ were imaged with a window of 3X zoom and 512×256 resolution was used for acquisition. Acquisition line speed was set to 8000 Hz with line average accumulation set to 1. 210 stacks were acquired bidirectionally using a Piezo focus kit MIPOS 500 with 6 planes per stack, and 10 µm distance between each plane. The acquisition rate was 0.813 Hz. 12 s baseline was acquired before stimulation, and each stimulation lasts for 12 s.

A capillary tube containing tastant was secured on a micromanipulator. To stimulate the fly proboscis, the position of the tube was adjusted to be perfectly surrounding the extended proboscis. Tastants were presented to the fly with ddH_2_O first with the rest in mixed orders, except for imaging dose-dependent responses which tastants were applied with ascending concentrations. Capillary was thoroughly washed with ddH_2_O before applying the next tastant.

### Imaging analysis

For TPN and LN 2-dimensional time series analysis, normalized fluorescence ΔF/F_0_ was calculated for selected ROIs that enclose the axonal projections. 10 representative points within the 10 s baseline were selected and averaged as F_0_. ΔF was calculated by dividing the mean fluorescence values by F_0_. Peak ΔF/F_0_ was calculated by averaging 3 points of the peak within the 5 s stimulation period. Due to the low baseline of GNG.SLP.T1 projections, a circular ROI with diameter of 30 µm was drawn. The edge of the circular ROI was 20 µm from the axon that traverses around the axonal projections. Statistical analyses were performed using Graphpad Prism 9.0.0, RStudio, and Python.

For volumetric imaging of ACh and GABA neurons in the SEZ, the acquired images were motion-corrected using CaImAn toolbox in Mesmerize (79, 80). Images were then aligned using MultiStackReg in ImageJ. For each experiment, all stimulation stacks were concatenated based on imaging planes using ImageJ. A calcium imaging computational toolbox by Romano et al. (2017) was used to check for imaging artifacts first and then segment single-neuron ROIs automatically based on morphological criteria (81). Automatically detected ROIs were checked and more ROIs were manually added or deleted if needed. To calculate ΔF/F_0_, 10 points of baseline before stimulation were averaged as F_0_. ΔF was calculated by dividing the mean fluorescence values by F_0_. Peak ΔF/F_0_ was calculated by averaging 3 points of the peak within the 12 s stimulation period. The ATP responses were calculated by deducting 200 mM NaCl ΔF/F_0_ responses from 100 mM 2Na·ATP ΔF/F_0_ responses.

To profile responses of ACh and GABA neurons in the SEZ, peak ΔF/F_0_ responses for stimulations of each ROI were binarized with a 10% response threshold. ROIs with all stimulation responses below 10% were treated as non-responders and excluded from later analysis. ROIs responding to only one tastant were treated as single responders whereas the rest were treated as multi-responders. Heatmaps, pie charts, response traces, bar graphs, and stacked-bar graphs were plotted with Seaborn 0.12.2 and Matplotlib 3.8.0 library in python.

### Connectomic analysis

Sweet/water (Gr64f/Ppk28), bitter (Gr66a), IR94e, and Ppk23^glut^ GRNs from left and right hemispheres were identified based on neuron projection morphology, predicted neurotransmitter expression (62), and public identification contributed by flywire community users (see credit table for details of contribution) on Codex.flywire.ai (v783, data acquired in February 2024). Neurons with unclear neurotransmitter prediction were excluded from analysis. Postsynpatic cholinergic and GABAergic LNs and putative directly connected TPNs were found by searching the downstream neurons of all the GRNs. Putative indirectly connected TPNs in the feedforward pathway were found by looking downstream of all the cholinergic LNs that are postsynaptic to all the GRNs. The number of synapses was recorded to examine the neural connectivity. To perform hierarchical clustering and UMAP visualization of the GRNs based on cholinergic LN connectivity, the matrix was first converted to a Compressed Sparse Row Matrix (csr matrix) and normalized using “l2” norm. TruncatedSVD was used to perform dimensional reduction to 10 components with a random state of 42. UMAP was plotted with random state = 42, n_neighbors = 40, min_dist = 0.2, and Euclidean metric. Hierarchical clustering was performed with “average” method and Euclidean distance metric. 11 clusters were chosen based on Silhouette score. Similar operations were done for the GRN vs GABAergic LN connectivity with several differences: n_neighbors was set to 10, and “ward” method was used for clustering. 8 clusters were chosen based on the Silhouette score. For GRN clustering based on directly connected TPN connectivity, dimensions were reduced to 10 components. Hierarchical clustering using “average” method and “correlation” metric was performed. 10 clusters were chosen based on Silhouette score. UMAP was performed with n_neighbors=100, min_dist=0.3, and “correlation” metric. For cholinergic LN clustering based on upstream Sweet/water GRN connectivity, dimensions were reduced to 10 components. Hierarchical clustering using “average” method and “correlation” metric was performed. 6 clusters were chosen based on Silhouette score. UMAP was performed with n_neighbors=30, min_dist=0.5, and “correlation” metric. For cholinergic LN clustering based on indirectly connected TPN connectivity, dimensions were reduced to 10 components. Hierarchical clustering using “ward” method and “euclidean” metric was performed. Pairwise cosine similarity, which calculates the similarity between two vectors in an inner product space, was used to compare the pairwise similarity of GRNs based on cholinergic LNs, GABAergic LNs, and putative directly connected TPNs connectivity.

### Computational modeling

Brain model experiments were performed using the leaky-integrate-and-fire model as detailed in Shiu et al., 2023 (60). Each class of GRNs were entered as input neurons, and 20 experiments with GRN firing rate ranging from 10-200 Hz with increments of 10 Hz were performed. The firing rates of all responding neurons in the whole brain were obtained. ACh interneurons, GABA interneurons, putative directly and indirectly connected TPNs were extracted from the list and their responses were profiled. The maximum activation level of each GRN class was determined based on the plateau point where the number of neurons activated no longer increases rapidly.

### NBLAST morphology analysis

To generate the average NBLAST score in Figure 7C for unshuffled and shuffled sets, all-by-all NBLAST similarity score of TPNs postsynaptic to each cholinergic LN was generated. We only focused on 37 cholinergic LNs that connect to 3 or more postsynaptic TPNs. Shuffling is within each cholinergic LN, meaning that if a cholinergic LN connects to 5 TPNs in the unshuffled dataset, 5 TPNs will be randomly selected from the pool of 151 TPNs. After generating the all-by-all comparison, the self-self comparison yielding a score of 1, and the duplicating scores from the same comparison were removed, and the remaining scores were averaged.

Hierarchical clustering on the TPNs was performed by generating all-by-all NBLAST comparison scores and calculating the distance metrics. Ward’s method was used for clustering and the dendrogram was cut at distance=4 to generate 4 clusters. To generate the probability distribution of shuffled set in Figure 7D, the probability of 2 postsynaptic TPNs belonging to the same cluster for individual cholinergic LNs was calculated as:

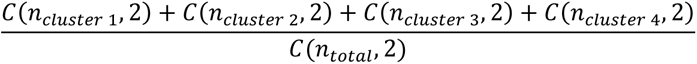

Where C indicates combination, n_total_ indicates the total number of TPNs postsynaptic to the given cholinergic LN, and n_cluster x_ indicates the number of postsynaptic TPNs belonging to cluster 1-4. 37 probability scores were generated and were plotted using descriptive statistics. For shuffled sets, the TPNs previously organized based on the clustering order were shuffled 1000 times, and each shuffle should assign each TPN with a random cluster identity. The same operation for calculating the probability was done for each shuffled set. To plot the histograms, the probabilities for each shuffled set were first binned and the average counts in each bin were plotted as the height of the histograms.

### Immunohistochemistry

Immunofluorescence staining for the *Drosophila* brain was performed similar to previous studies (40, 41). Briefly, flies were fixed in 4% paraformaldehyde in PBST. Fly brains were dissected and incubated in primary antibodies (rabbit anti-GFP in 1:1000 and mouse anti-brp in 1:50) for 24-48 hours. The brains were washed in 0.1% PBST and secondary antibodies (goat anti-rabbit Alexa 488 in 1:200 and goat anti-mouse Alexa 647 in 1:200) were added. The immunofluorescence stacks were acquired with a Leica SP5 II Confocal under a 25x water immersive objective. The stacks were later processed in imageJ.

## Acknowledgments

We thank the Princeton FlyWire team and members of the Murthy and Seung labs, as well as members of the Allen Institute for Brain Science, for development and maintenance of FlyWire (supported by BRAIN Initiative grants MH117815 and NS126935 to Murthy and Seung). We also acknowledge members of the Princeton FlyWire team and the FlyWire consortium for neuron proofreading and annotation. We thank Bloomington Drosophila Stock Center for the transgenic flies. We thank Dr. Mark Cembrowski for guidance on the clustering analyses. We thank the Gordon Lab members for valuable advice on the project. Funding for this project was from Canadian Institutes of Health Research (CIHR) grants FDN-148424 and NSERC Discovery Grant RGPIN-2016-03857.

## Author Contributions

J.L. and M.D.G. developed the project. M.D.G acquired funding. J.L. and R.D. performed connectomics analyses. M.S. performed preliminary calcium imaging on marge neurons. J.L. performed calcium imaging and brain model experiments. P.J. identified split-Gal4 lines. J.L. and M.D.G. wrote the manuscript.

## Code availability

https://github.com/jil713/Li-et-al-2024-connectome

## Raw data availability

https://data.mendeley.com/preview/cnxyw4sykb?a=bad193d2-8423-4c2f-9bce-6794649f90e3

