## Supplemental Figures for "Functional imaging and connectome analyses reveal organizing principles of taste circuits in *Drosophila*"

**Supplementary Information**

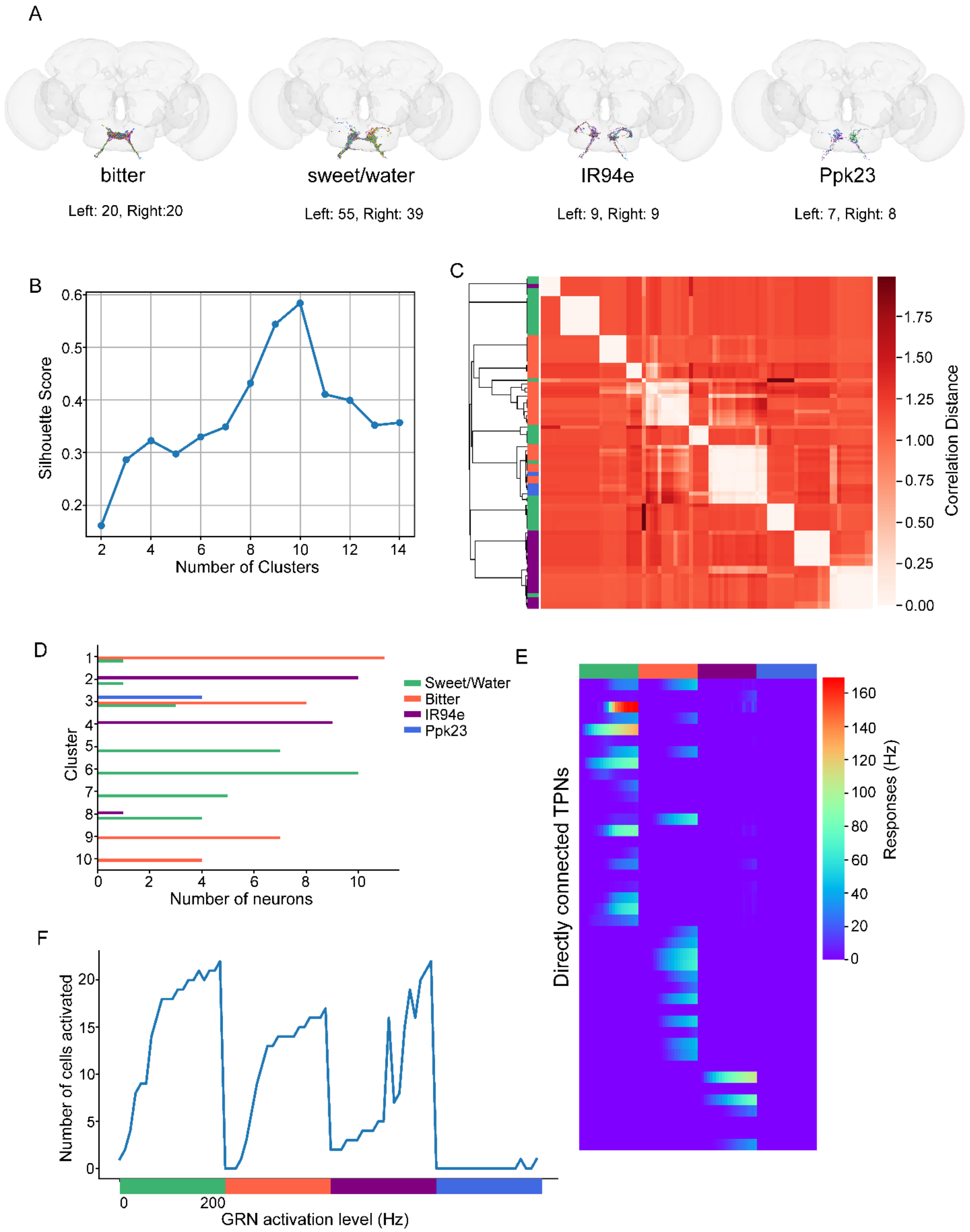

**Figure S1: GRN identification, cluster analysis detail, and detailed brain model results. Related to Figure 1.**

**A)** Neuron skeletons of the identified GRN types in the connectome. **B)** Average Silhouette score for 2-14 clusters. 10 clusters were chosen based on maximum separation indicated by the peak of the Silhouette score. **C)** Pairwise correlation distance between each GRN in the hierarchical clustering based on connectivity with TPNs. **D)** Detailed composition of GRN types within each cluster. **E)** Response heatmap of directly connected TPN responses to 10-200 Hz stimulation of each GRN type in the brain model. Solid colored bars indicate GRN type activated at increasing frequencies from left to right **F)** The number of directly connected TPNs activated with 10-200 Hz stimulation of each GRN type. Graph was used to determine the appropriate maximum activation level for each modeling experiment.

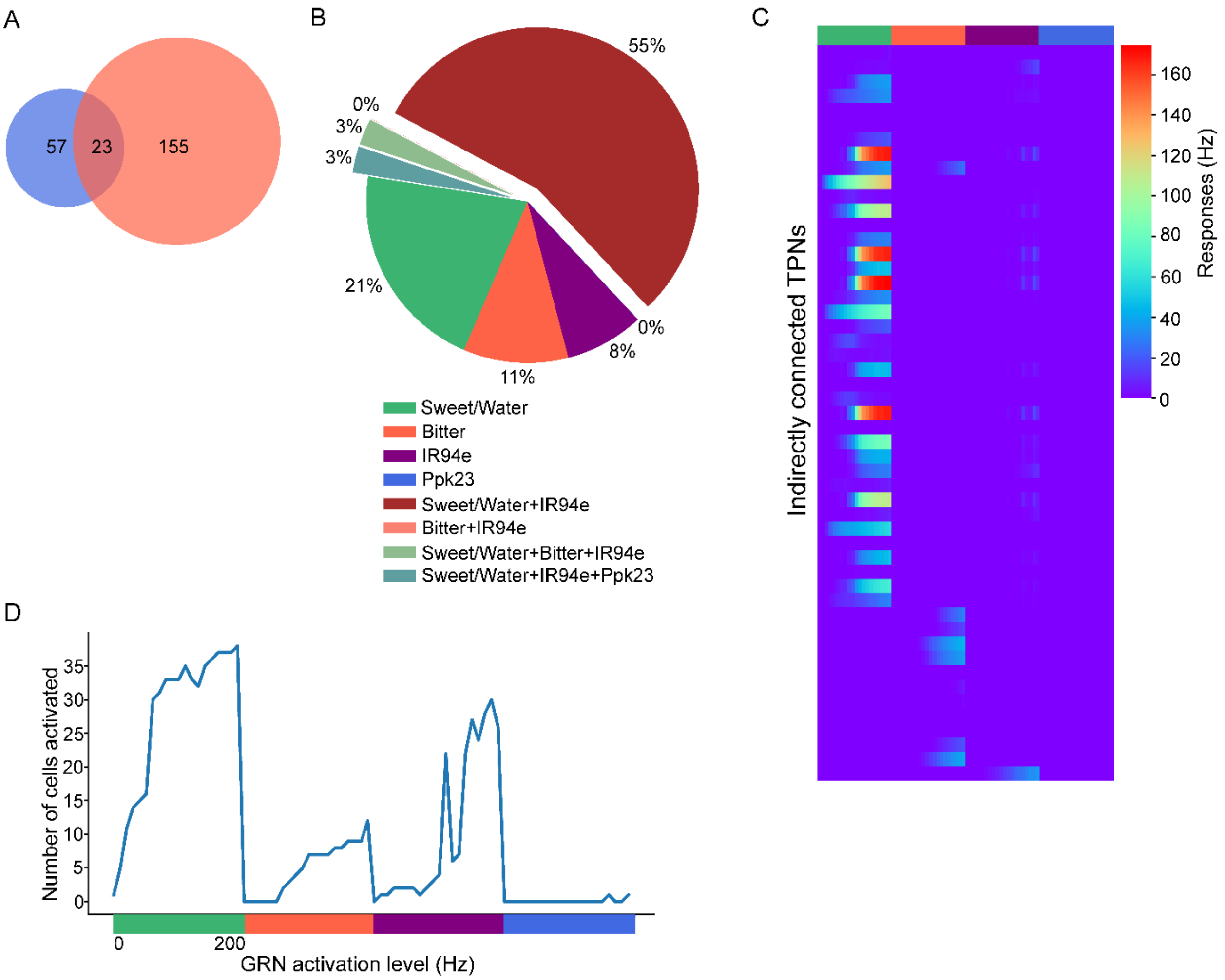


**Figure S2: Number of identified indirectly connected TPNs from the connectome and tuning properties based on the brain model. Related to Figure 1.**

**A)** Relationship between directly and indirectly connected TPNs. 57 TPNs are directly connected to GRNs, and 155 TPNs are indirectly connected, with 23 neurons shared between the two populations. **B)** Proportion of indirectly connected TPNs displaying each tuning profile in the brain model. **C)** Response heatmap of indirectly connected TPNs following activation of different GRN types at levels of 10-200 Hz in the brain model. **D)** The number of indirectly connected TPNs activated with 10-200 Hz stimulation of each GRN type.

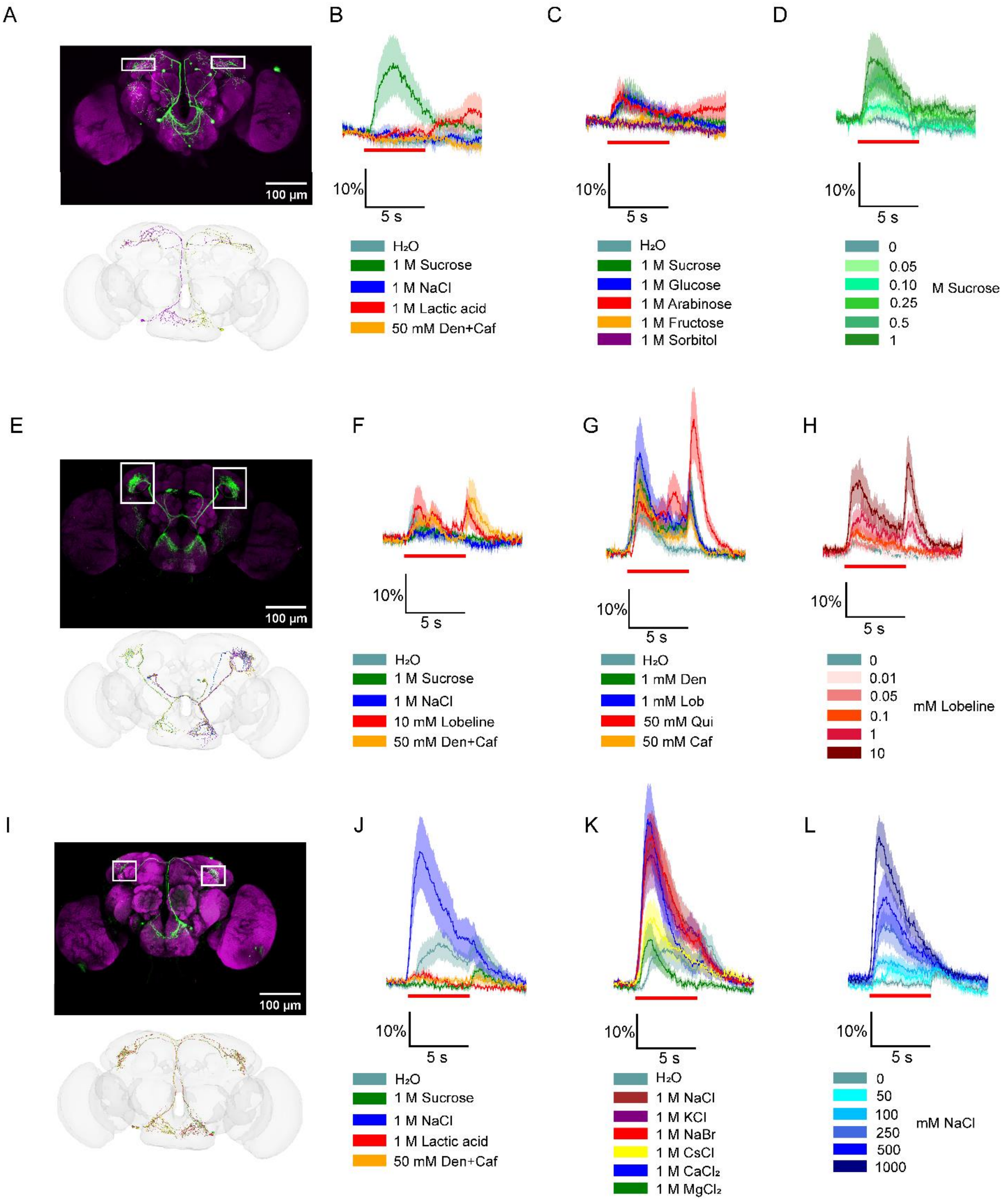


**Figure S3: Calcium imaging traces of example TPNs. Related to Figure 2.**

**A)** Immunostaining (top) and connectome neuron skeleton (bottom) of peafowl. **B-D)** Calcium traces of peafowl in response to different taste qualities (B), sugars (C), and concentrations of sucrose (D). **E)** Immunostaining (top) and connectome neuron skeleton (bottom) of mlSEZt. **F-G)** Calcium traces of mlSEZt in response to different taste qualities (F), bitters (G), and concentrations of lobeline (H). **I)** Immunostaining (top) and connectome neuron skeleton (bottom) of GNG.SLP.T1. **J-L)** Calcium traces of GNG.SLP.T1 in response to different taste qualities, salts, and concentrations of NaCl, respectively. Red boxes on immunostained images indicate area imaged during calcium imaging.

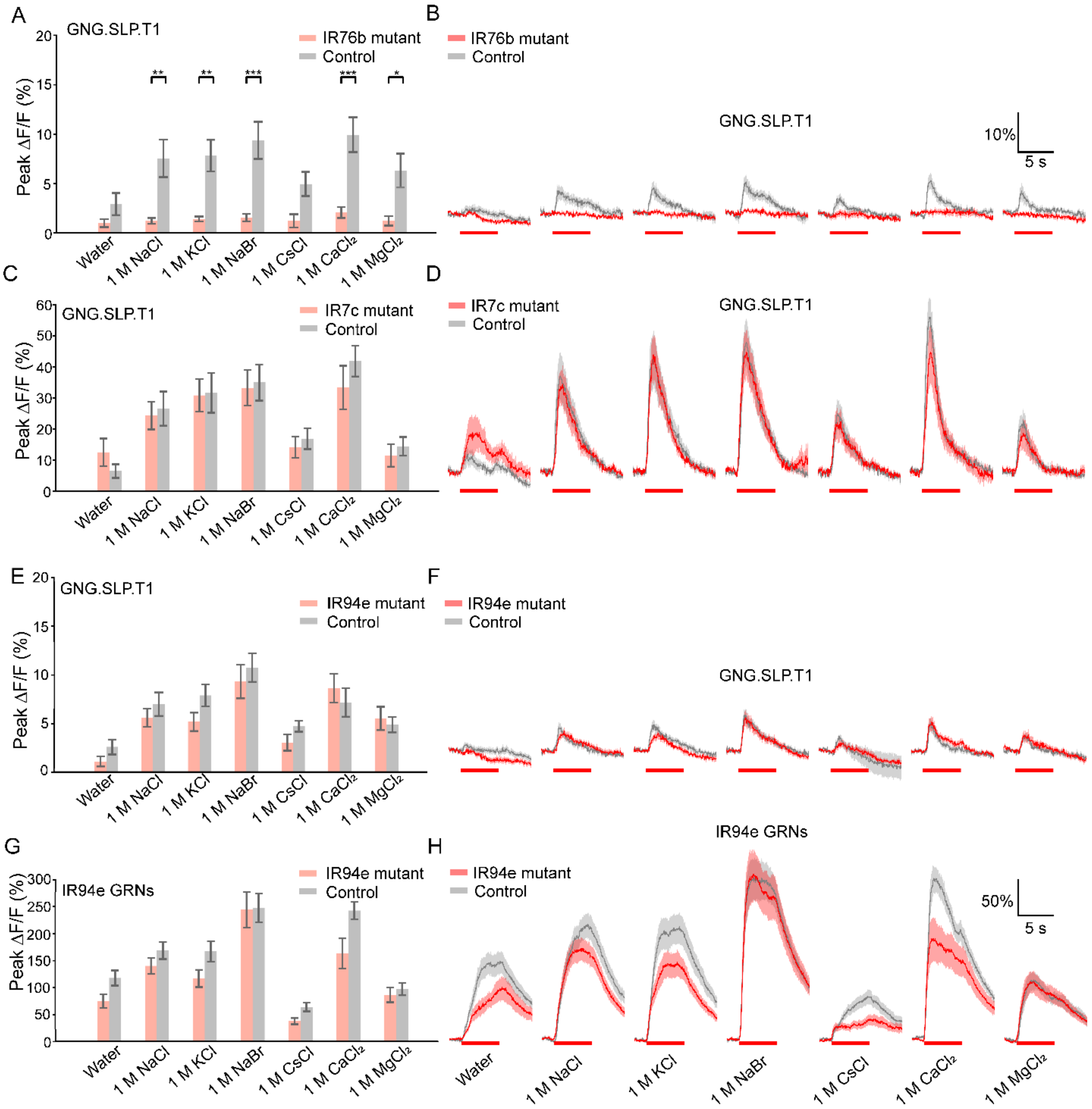


**Figure S4: Calcium imaging traces of GNG.SLP.T1 TPNs with different mutants. Related to Figure 2.**

**A-B)** Peak calcium responses (A) and response traces (B) of GNG.SLP.T1 in *IR76b* mutants stimulated by various monovalent and divalent salts. **C-D)** Peak calcium responses (C) and response traces (D) of GNG.SLP.T1 in *IR7c* mutants stimulated by various monovalent and divalent salts. **E-F)** Peak calcium responses (E) and response traces (F) of GNG.SLP.T1 in *IR94e* mutants stimulated by various monovalent and divalent salts. **G-H)** Peak calcium responses (G) and response traces (H) of IR94e sensory neurons in *IR94e* mutants stimulated by various monovalent and divalent salts.

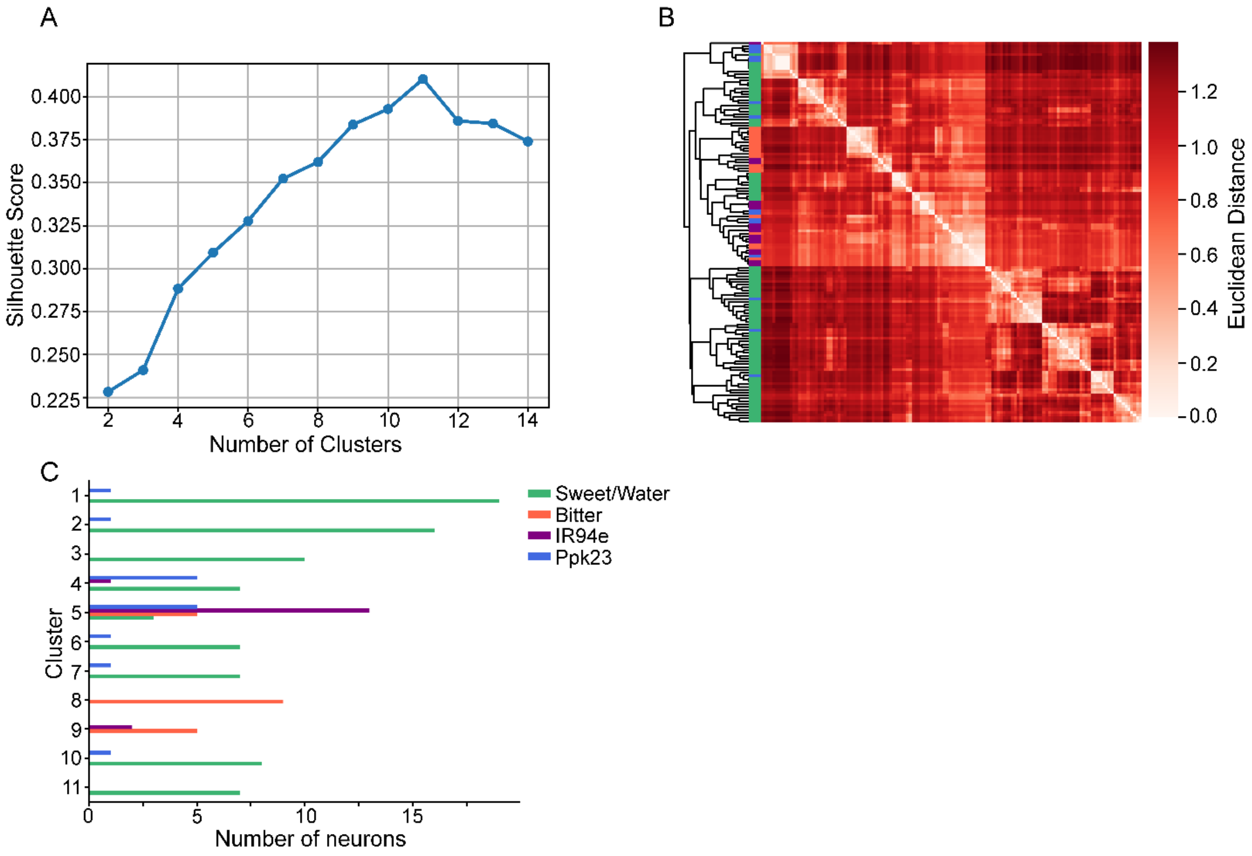


**Figure S5: Details of GRN clustering based on cholinergic LN connectivity. Related to Figure 3.**

**A)** Average silhouette score for 2-14 clusters. 11 clusters were chosen based on the peak silhouette score. **B)** Pairwise Euclidean distance from the hierarchical clustering of the GRNs based on cholinergic LN connectivity. **C)** Detailed composition of GRNs in each cluster.

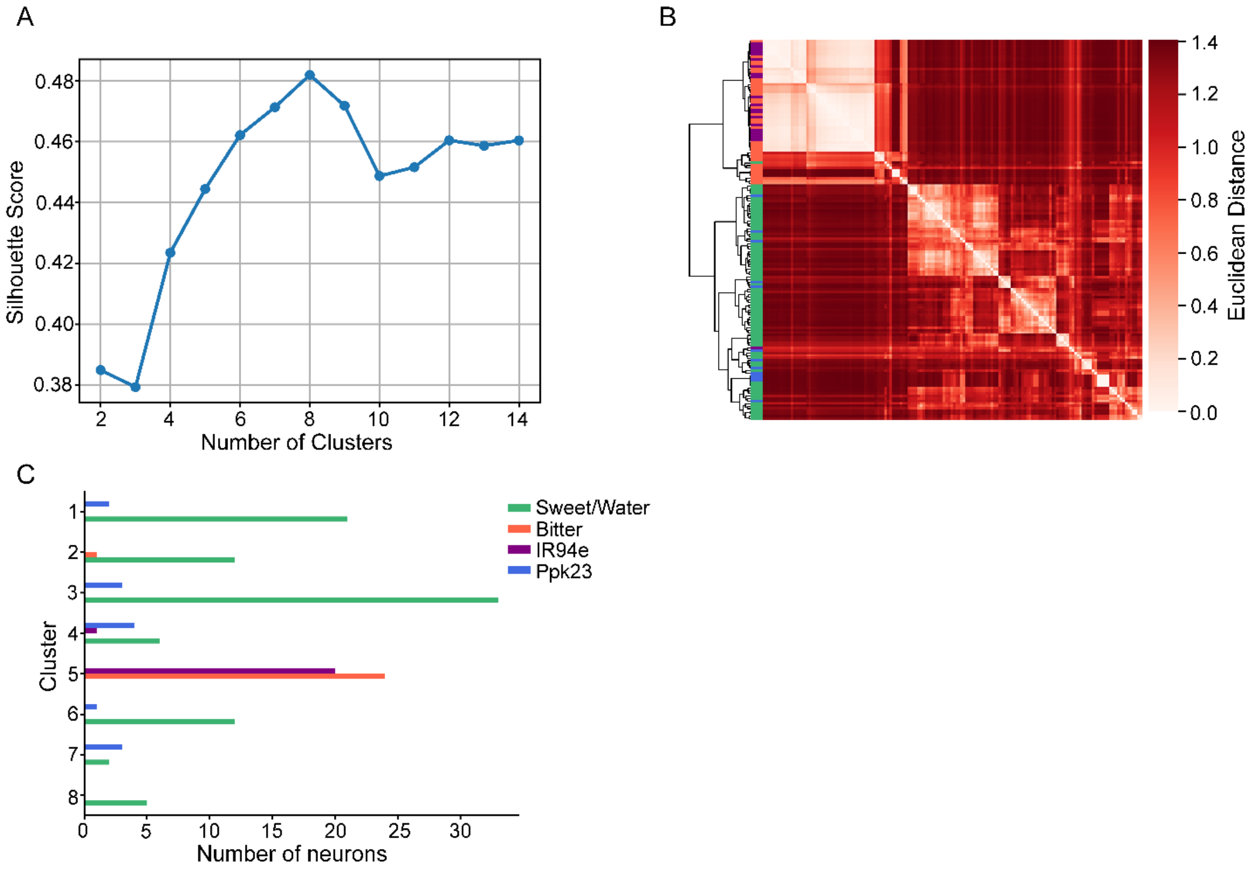


**Figure S6: Details of GRN clustering based on GABAergic LNs. Related to Figure 4.**

**A)** Average silhouette score for 2-14 clusters. 8 clusters were determined based on the peak silhouette score. **B)** Pairwise Euclidean distance from the hierarchical clustering of GRNs based on GABAergic LN connectivity. **C)** Detailed composition of GRNs in each cluster.

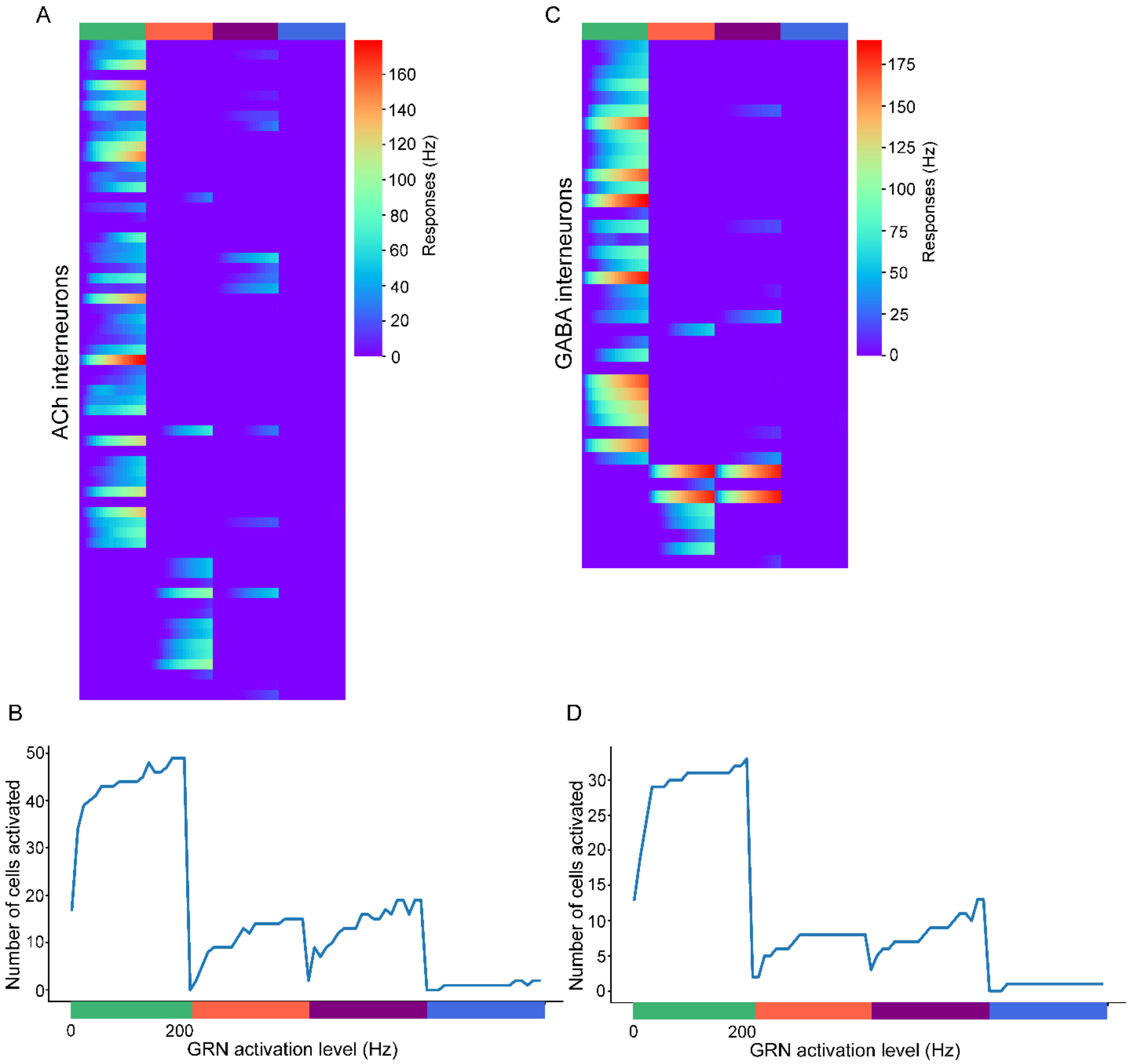


**Figure S7: Detailed activation levels of cholinergic and GABAergic LNs in the brain model. Related to Figure 5.**

**A)** Heatmap of cholinergic LN responses to different GRN type stimulation at levels 10-200 Hz in the brain model. **B)** The number of cholinergic LNs activated with 10-200 Hz stimulation of each GRN type. **C)** Heatmap of GABAergic LN responses to different GRN type stimulation at levels 10-200 Hz in the brain model. **D)** The number of GABAergic LNs activated with 10-200 Hz stimulation of each GRN type.

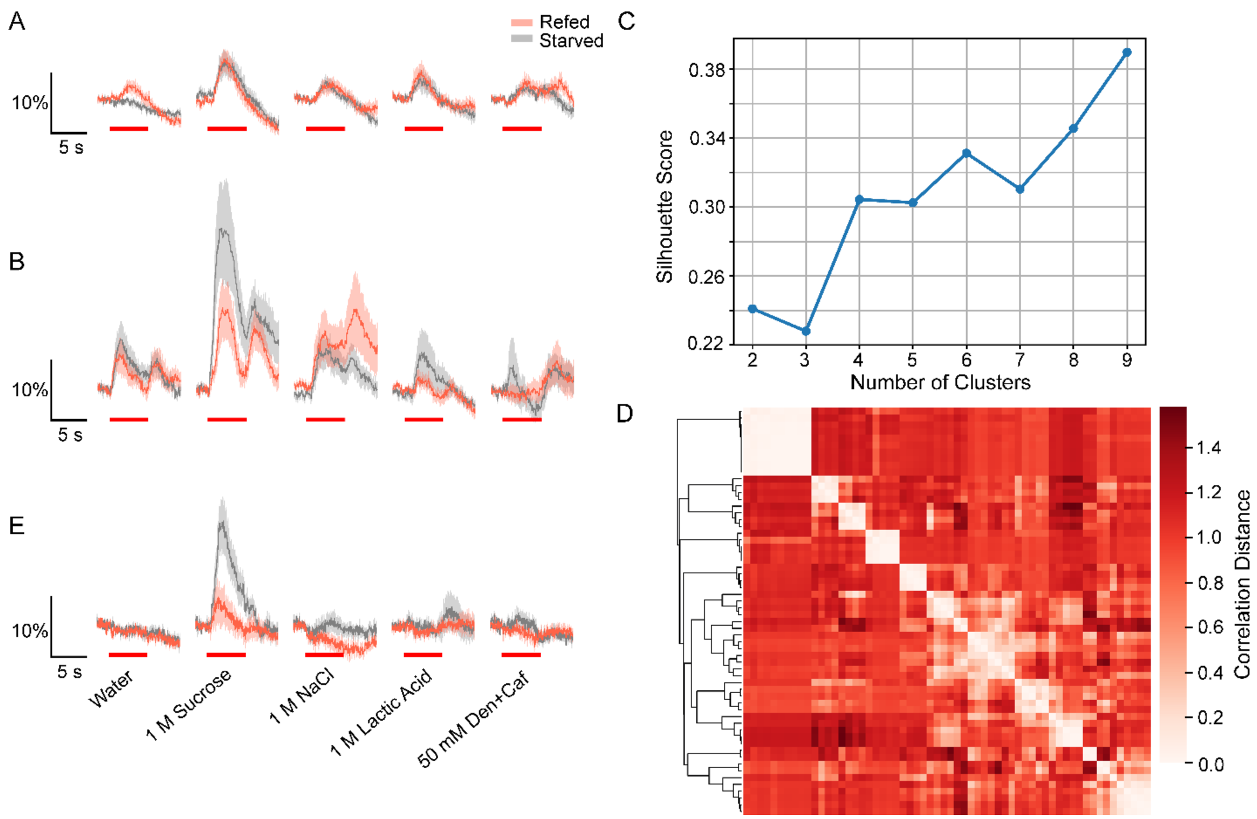


**Figure S8: Calcium imaging traces of selected cholinergic LNs, and clustering details of cholinergic LNs that receive sweet GRN inputs. Related to Figure 6.**

**A)** Calcium response traces of rattle LNs in response to different taste qualities under refed and starved conditions. **B)** Calcium response traces of marge LNs in response to different taste qualities under refed and starved conditions. **C)** Average silhouette score for 2-9 cholinergic LN clusters. 6 clusters were chosen based on the peak silhouette score within plausible number of clusters. **D)** Pairwise correlation distance matrix from the hierarchical clustering of the cholinergic LNs postsynaptic to sweet/water GRNs. **E)** Calcium traces of peafowl TPNs, which are postsynaptic to marge LNs, in response to different taste qualities under refed and starved conditions.

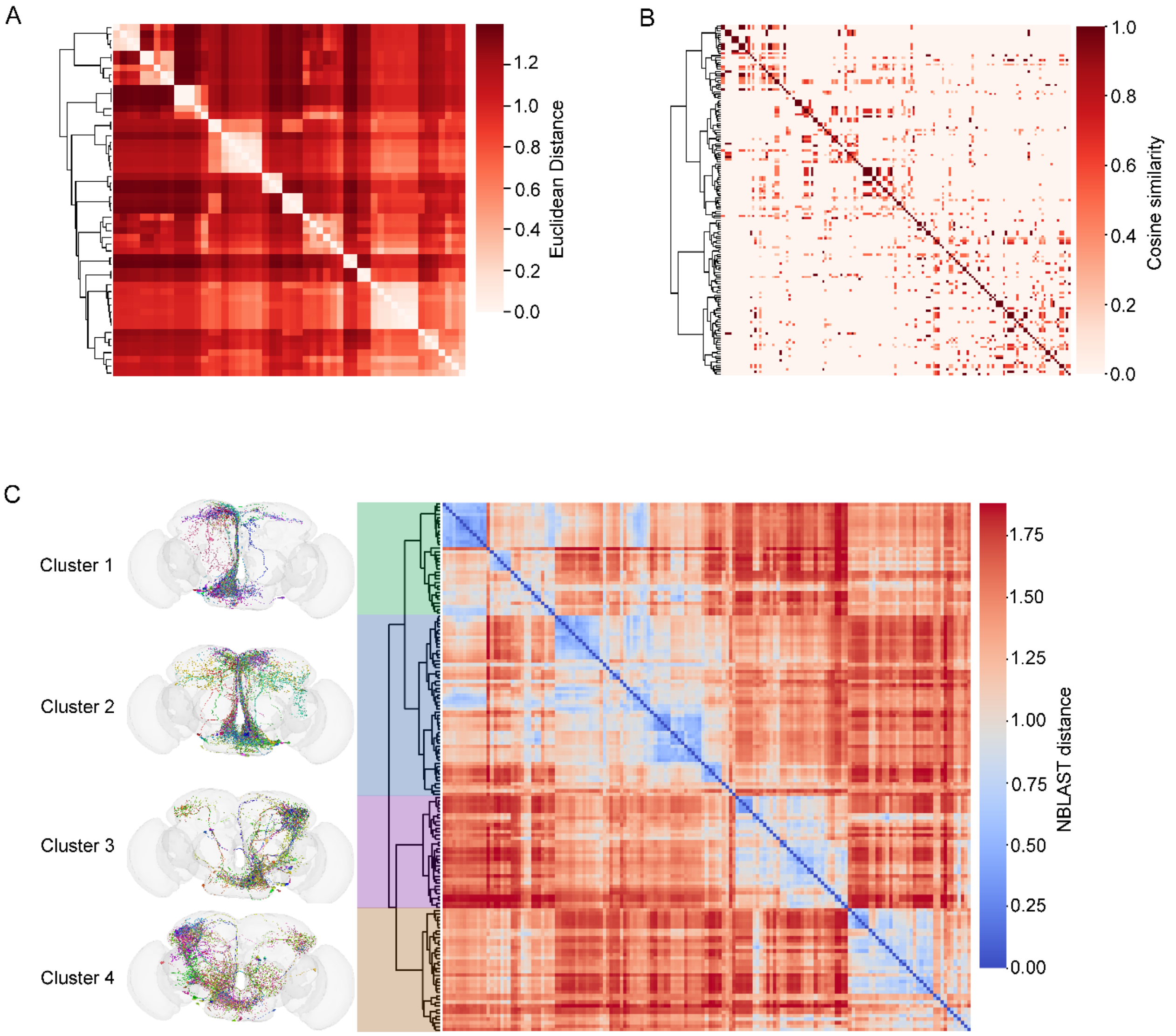


**Figure S9: Morphological comparison of the third-order TPNs. Related to Figure 7.**

**A)** Pairwise Euclidean distance matrix from the hierarchical clustering of the cholinergic LNs based on postsynaptic TPN connectivity. **B)** Pairwise cosine similarity of connectivity between pairs of TPNs organized based on morphological clustering. **C)** Pairwise NBLAST distance of morphological clustering of all the indirectly connected TPNs. The dendrogram was cut at distance of 4 to generate 4 clusters of TPNs based on morphology.

**Key Resources Table**

**Table S1: Reagents used in the study.**

| **Reagent or Resource** | **Source** | **Identifier** |
|---|---|---|
| **Chemicals** | | |
| D-Sucrose | Sigma-Aldrich | S7903 |
| D-Fructose | Sigma-Aldrich | F0127 |
| D-Glucose | Sigma-Aldrich | G5400 |
| D-Arabinose | Sigma-Aldrich | A3131 |
| D-Sorbitol | Sigma-Aldrich | S1876 |
| Denatonium benzoate | Sigma-Aldrich | D5765 |
| Quinine hydrochloride dihydrate | Sigma-Aldrich | Q1125 |
| Caffeine | Sigma-Aldrich | C0750 |
| Lobeline hydrochloride | Sigma-Aldrich | 141879 |
| DL-Lactic acid | Sigma-Aldrich | 69785 |
| NaCl | Sigma-Aldrich | S7653 |
| KCl | Sigma-Aldrich | P9541 |
| NaBr | Sigma-Aldrich | S4547 |
| CsCl | Sigma-Aldrich | 289329 |
| CaCl2 | BDH chemicals | BDH4524 |
| MgCl2 | BDH chemicals | BDH0244 |
| 2Na·ATP | Sigma-Aldrich | A1852 |
| Polyethylene glycol (PEG) | Sigma-Aldrich | 202444 |
| Agar | Sigma-Aldrich | A1296 |
| 4% paraformaldehyde in PBS | Thermo Fisher Scientific | J61899.AK |
| Rabbit anti-GFP | Invitrogen | A11122, RRID: AB_221569 |
| Mouse anti-brp (nc82) | DSHB | RRID: AB_2392664 |
| Alexa Fluor 488 goat anti-rabbit IgG | Invitrogen | A11008, RRID: AB_143165 |
| Alexa Fluor 647 goat anti-mouse IgG | Invitrogen | A11126, RRID: AB_2535804 |
| **Fly strains** | | |
| *D. melanogaster: w ; P{20XUAS-IVS-GCaMP6m}attP40* | Bloomington Drosophila Stock Center | BDSC: 42748 |
| *D. melanogaster*: w ; PBac{20XUAS-IVS-GCaMP6m}VK00005 | Bloomington Drosophila Stock Center | BDSC: 42750 |
| *D. melanogaster: w ; P{20XUAS-IVS-GCaMP6f}attP40* | Bloomington Drosophila Stock Center | BDSC: 42747 |
| *D. melanogaster: w[1118]; P{y[+t7.7] w[+mC]=20XUAS-IVS-jGCaMP7s}su(Hw)attP5* | Bloomington Drosophila Stock Center | BDSC: 80905 |
| *D. melanogaster: w; P{20XUAS-IVS-GCaMP6s}attP40* | Bloomington Drosophila Stock Center | BDSC: 42746 |
| *D. melanogaster: Peafowl* | Sterne et al. (2021)(1) | Janelia Stock Center: SS46596 |

| | | |
|---|---|---|
| *D. melanogaster: R43D01-AD* | Bloomington Drosophila Stock Center | BDSC: 70690 |
| *D. melanogaster: R29F12-DBD* | Bloomington Drosophila Stock Center | BDSC: 68893 |
| *D. melanogaster: VT019729-AD* | Bloomington Drosophila Stock Center | BDSC: 73020 |
| *D. melanogaster: VT008484-DBD* | Bloomington Drosophila Stock Center | BDSC: 74557 |
| *D. melanogaster:* $IR76b^1$ | Zhang et al. (2013) (2) | Flybase: FBst0051309 |
| *D. melanogaster:* $IR76b^2$ | Zhang et al. (2013) (2) | Flybase: FBst0051310 |
| *D. melanogaster:* $IR7cGal4^{RFP}$ | McDowell et al. (2022) (3) | |
| *D. melanogaster:* $IR94eLexA^{RFP}$ | McDowell et al. (2022) (3) | Flybase: FBal0376356 |
| *D. melanogaster: IR94e-Gal4* | Tirian and Dickson (2017) (4) | VDRC: v207582 |
| *D. melanogaster: Cha-Gal4* | Salvaterra and Kitamoto (2001) (5) | |
| *D. melanogaster: Gad1-Gal4* | Bloomington Drosophila Stock Center | BDSC: 51630 |
| *D. melanogaster: R18H03-AD* | Bloomington Drosophila Stock Center | BDSC: 69804 |
| *D. melanogaster: VT046049-DBD* | Bloomington Drosophila Stock Center | BDSC: 75523 |
| *D. melanogaster: Rattle* | Sterne et al. (2021), Shiu et al. (2022)(1, 6) | Janelia Stock Center: SS46917 |

**Table S2: Detailed fly genotypes used in the study.**

| Figure panel | Genotype |
|---|---|
| Figure 2A-D, Figure S3B-D | w; UAS-GCaMP6m/VT008469-AD; UAS-GCaMP6m/VT000926-DBD |
| Figure 2E-H, Figure S3F-H | w; UAS-GCaMP6f/R43D01-AD; +/R29F09-DBD |
| Figure 2I-L, Figure S3J-L | w; UAS-GCaMP6m/VT019729-AD; UAS-GCaMP6m/VT008484-DBD |
| Figure S4A-B | w; UAS-jGCaMP7s/VT019729-AD; IR76b$^{1}$/IR76b$^{2}$, VT008484-DBD |
| | w; UAS-jGCaMP7s/VT019729-AD; +/VT008484-DBD |
| Figure S4C-D | IR7cGAL4$^{RFP}$/IR7cGAL4$^{RFP}$; UAS-GCaMP6m/VT019729-AD; UAS-GCaMP6m/VT008484-DBD |
| | w; UAS-GCaMP6m/VT019729-AD; UAS-GCaMP6m/VT008484-DBD |
| Figure S4E-F | w; UAS-jGCaMP7s/VT019729-AD; IR94eLexA$^{RFP}$/IR94eLexA$^{RFP}$, VT008484-DBD |
| | w; UAS-jGCaMP7s/VT019729-AD; +/VT008484-DBD |
| Figure S4G-H | w; UAS-jGCaMP7s/Bl; IR94eLexA$^{RFP}$/IR94eLexA$^{RFP}$, IR94e-Gal4 |
| | w; UAS-jGCaMP7s/+; IR94e-Gal4/TM2 |
| Figure 5A-D, Figure S7A | w; UAS-GCaMP6s/Cha-Gal4; IR94eLexA$^{RFP}$/LexAop-P2X$_2$ |
| Figure 5E-H, Figure S7B | w; UAS-GCaMP6s/Gad1-Gal4; IR94eLexA$^{RFP}$/LexAop-P2X$_2$ |
| Figure 6B, Figure S9A | w; UAS-GCaMP6m/R81E10-AD; UAS-GCaMP6m/VT023745-DBD |
| Figure 6D, Figure S9B | w; UAS-GCaMP6m/R18H03-AD; +/VT046049-DBD |
| Figure 6H, Figure S9E | w; UAS-GCaMP6m/VT008469-AD; UAS-GCaMP6m/VT000926-DBD |
